# Bidirectional genome-wide CRISPR screens reveal host factors regulating SARS-CoV-2, MERS-CoV and seasonal coronaviruses

**DOI:** 10.1101/2021.05.19.444823

**Authors:** Antoine Rebendenne, Priyanka Roy, Boris Bonaventure, Ana Luiza Chaves Valadão, Lowiese Desmarets, Yves Rouillé, Marine Tauziet, Mary Arnaud-Arnould, Donatella Giovannini, Yenarae Lee, Peter DeWeirdt, Mudra Hegde, Francisco Garcia de Gracia, Joe McKellar, Mélanie Wencker, Jean Dubuisson, Sandrine Belouzard, Olivier Moncorgé, John G. Doench, Caroline Goujon

## Abstract

Several genome-wide CRISPR knockout screens have been conducted to identify host factors regulating SARS-CoV-2 replication, but the models used have often relied on overexpression of ACE2 receptor. Additionally, such screens have yet to identify the protease TMPRSS2, known to be important for viral entry at the plasma membrane. Here, we conducted a meta-analysis of these screens and showed a high level of cell-type specificity of the identified hits, arguing for the necessity of additional models to uncover the full landscape of SARS-CoV-2 host factors. We performed genome-wide knockout and activation CRISPR screens in Calu-3 lung epithelial cells, as well as knockout screens in Caco-2 intestinal cells. In addition to identifying ACE2 and TMPRSS2 as top hits, our study reveals a series of so far unidentified and critical host-dependency factors, including the Adaptins AP1G1 and AP1B1 and the flippase ATP8B1. Moreover, new anti-SARS-CoV-2 proteins with potent activity, including several membrane-associated Mucins, IL6R, and CD44 were identified. We further observed that these genes mostly acted at the critical step of viral entry, with the notable exception of ATP8B1, the knockout of which prevented late stages of viral replication. Exploring the pro- and anti-viral breadth of these genes using highly pathogenic MERS-CoV, seasonal HCoV-NL63 and -229E and influenza A orthomyxovirus, we reveal that some genes such as AP1G1 and ATP8B1 are general coronavirus cofactors. In contrast, Mucins recapitulated their known role as a general antiviral defense mechanism. These results demonstrate the value of considering multiple cell models and perturbational modalities for understanding SARS-CoV-2 replication and provide a list of potential new targets for therapeutic interventions.

## Introduction

Severe acute respiratory syndrome coronavirus 2 (SARS-CoV-2) is the etiologic agent of the coronavirus disease 2019 (COVID-19) pandemic, which has been applying an unprecedented pressure on health systems worldwide since it was first detected in China at the end of 2019. As of today (May 18, 2021), SARS-CoV-2 continues to spread worldwide, with over 164 million confirmed cases and >3,4 million deaths.

SARS-CoV-2 is the third highly pathogenic coronavirus to cross the species barrier in the 21^st^ century and cause an epidemic in the human population after SARS-CoV(-1) in 2002-2003 (Drosten et al., 2003; Peiris et al., 2003; Zhong et al., 2003) and Middle East respiratory syndrome (MERS)-CoV in 2012 (Zaki et al., 2012). These three coronaviruses share some common clinical features, including breathing difficulty, acute respiratory distress syndrome (ARDS) and death in the most extreme cases (Zhou et al., 2020a). Four additional Human Coronaviruses (HCoV-229E, -NL63, -OC43 and -HKU1) are known to circulate seasonally in humans and are associated with multiple respiratory diseases of varying severity including common cold, pneumonia and bronchitis, contributing to approximately one-third of common cold infections in humans (van der Hoek, 2007).

Coronaviruses are enveloped, positive stranded RNA viruses with a genome of approximately 30 kilobases. Highly pathogenic SARS-CoV-1, MERS-CoV and SARS-CoV-2, as well as seasonal HCoV-OC43 and HCoV-HKU1, belong to the genus *betacoronavirus,* whereas seasonal HCoV-229E and HCoV-NL63 are alphacoronaviruses. The respiratory tract is the main replication site of SARS-CoV-2, but it has also been shown to replicate in the gastrointestinal tract (Xiao et al., 2020) and infect other cell types. Like SARS-CoV-1 and HCoV-NL63, SARS-CoV-2 entry into target cells is mediated by the Angiotensin converting enzyme 2 (ACE2) receptor (Hoffmann et al., 2020; Hofmann et al., 2005; Li et al., 2003; Wu et al., 2009; Zhou et al., 2020b). The cellular serine protease Transmembrane protease, serine 2 (TMPRSS2) is employed by both SARS-CoV-1 and -2 for Spike (S) protein priming at the plasma membrane (Hoffmann et al., 2020; Matsuyama et al., 2010). Cathepsins are also involved in SARS-CoV S protein cleavage and fusion peptide exposure upon entry via an endocytic route in the absence of TMPRSS2 (Huang et al., 2006; Ou et al., 2020; Simmons et al., 2005). Conversely, HCoV-229E entry into target cells is mediated by membrane aminopeptidase N (ANPEP) (Yeager et al., 1992), whereas MERS-CoV enters via dipeptidyl peptidase 4 (DPP4) (Raj et al., 2013). Importantly, both these coronaviruses are also known to use TMPRSS2 for S protein activation (Bertram et al., 2013; Gierer et al., 2013).

Following viral entry and delivery of the viral genomic RNA associated with the nucleocapsid (N) to the cytoplasm, ORF1a/b is directly accessible to the translation machinery, which leads to the synthesis of two polyproteins (pp), pp1a and pp1b. These polyproteins are further processed into nonstructural proteins, which are important for the formation of replication and transcription complexes. The replication/transcription steps take place at the endoplasmic reticulum (ER) through the active formation of replication organelles surrounded by double membranes, which form a protective microenvironment against viral sensors and restriction factors. Subgenomic RNAs are then transcribed, translated into structural proteins, and translocated to the ER. The assembly takes place at the endoplasmic reticulum-Golgi intermediate compartments, where newly produced genomic RNAs associated with N are also recruited. Budding occurs at the Golgi compartment and newly generated virions are released by exocytosis (reviewed in (V’kovski et al., 2020).

Coronaviruses, which are found throughout the animal kingdom with an important diversity in bats, have a particularly high potential for cross-species transmission and may be the origin of future pandemics (Irving et al., 2021). There is therefore a dire need to study coronaviruses in depth and to identify new therapeutic targets against these viruses.

Several whole-genome KO CRISPR screens for the identification of coronavirus regulators have been recently reported (Baggen et al., 2021; Daniloski et al., 2021; Schneider et al., 2021; Wang et al., 2021; Wei et al., 2021; Zhu et al., 2021). These screens used simian Vero E6 cells (Wei et al., 2021), human Huh7 cells (or derivatives) ectopically expressing ACE2 and TMPRSS2 or not (Baggen et al., 2021; Schneider et al., 2021; Wang et al., 2021), and A549 cells ectopically expressing ACE2 (Daniloski et al., 2021; Zhu et al., 2021). Here, we conducted genome-wide loss-of-function screens by CRISPR knockout (KO) and gain-of-function screens by CRISPR activation to identify host factors modulating SARS-CoV-2 infection. Naturally permissive simian Vero E6 cells, as well as physiologically relevant human lung epithelial Calu-3 cells and intestinal Caco-2 cells, were used in these screens. Well-known SARS-CoV-2 host dependency factors were identified as top hits, such as ACE2, and either TMPRSS2 or Cathepsin L (depending on the cell type), validating the rationale of this study. Moreover, ACE2 scored as the top enriched and top depleted hit in all CRISPR KO and activation screens in Calu-3 cells, respectively, underlying the complementarity of both approaches. We validated the role of our top hits using individual CRISPR KO or activation in Calu-3 cells and assessed their effect on other coronaviruses and orthomyxovirus influenza A. Altogether, this quantitative and integrative study provides new insights in SARS-CoV-2 life cycle by identifying new host factors that modulate either positively or negatively replication of SARS-CoV-2 and other coronaviruses, and might lead to new, pan-coronavirus strategies for host-directed therapies.

## Results

### Meta-analysis of existing CRISPR KO screen data highlights the importance of diverse models

African Green Monkey (*Chlorocebus sabaeus*) Vero E6 cells of kidney origin are commonly used to amplify SARS-CoV-2 and present high levels of cytopathic effects (CPE) upon replication, making them ideal to perform whole-genome CRISPR screens for host factor identification. A *C. sabaeus* sgRNA library was previously described and successfully used to identify host factors regulating SARS-CoV-2 (isolate USA-WA1/2020) and other coronavirus replication (Wei et al., 2021). In order to determine whether hit identification based on whole-genome CRISPR screens was reproducible across different laboratories and virus isolates, we initially repeated whole-genome CRISPR KO screens in Vero E6 cells using the SARS-CoV-2 isolate BetaCoV/France/IDF0372/2020. Vero E6 cells were first stably engineered to express Cas9 and activity was validated with a GFP activity assay. We then transduced the cells with the *C. sabaeus* genome-wide pooled CRISPR library (Wei et al., 2021) at a low MOI (∼0.1-0.5) in biological duplicates, the first of which was then divided into three technical replicates. Cells were either collected for subsequent genomic DNA extraction or challenged with SARS-CoV-2 at MOI 0.005, a viral input that induced cell death in >95% of the cells in 3-4 days. After viral challenge, surviving cells were propagated for 11-13 days to increase cell numbers prior to genomic DNA extraction, PCR, and Illumina sequencing (**Figure 1A**). We determined the log2-fold-change (LFC) of guides, comparing SARS-CoV-2-challenged to untreated cells, and observed that replicates for these screens were well-correlated (**Figure S1A**)

**Figure 1.**
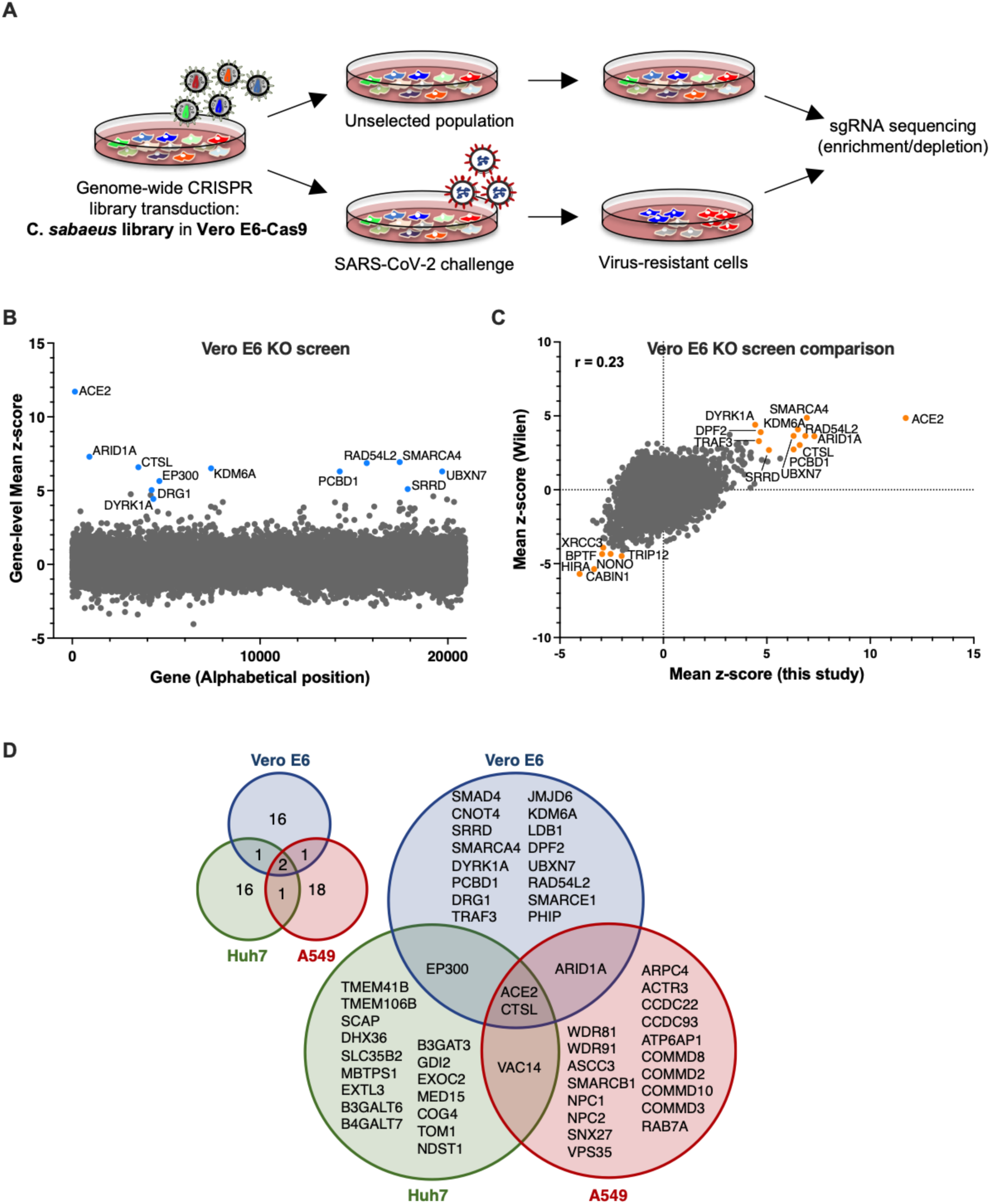
**Cell-type specificity of SARS-CoV-2 regulators identified by CRISPR screens.** **A.** Schematic of pooled screen to identify SARS-CoV-2 regulators in Vero E6 cells. **B.** Scatter plot showing the gene-level mean z-scores of genes when knocked out in Vero E6 cells. The top genes conferring resistance to SARS-CoV-2 are annotated and shown in blue. **C.** Comparison between this Vero E6 screen to the Vero E6 screen conducted by the Wilen lab (Wei et al., 2021). Genes that scored among the top 20 resistance hits and sensitization hits in both screens are labeled. **D.** Venn diagram comparing hits across screens conducted in Vero E6, A549, and Huh7 (or derivatives) cells (ectopically expressing ACE2 and TMPRSS2 or not). The top 20 genes from each cell line are included, with genes considered a hit in another cell line if the average Z-score > 3.

We first examined the results from this screen and saw that ACE2 was a top hit, among other genes (**Figure 1B**). Compared to the prior results from the Wilen lab, this screen showed greater statistical significance for pro-viral (resistance) hits, indicating that the screening conditions employed here resulted in stronger selective pressure (**Figure 1C**, **Figure S1B**). Nevertheless, pro-viral hits were consistent across the two screens, with 11 genes scoring in the top 20 of both datasets, including ACE2 and CTSL; similarly, 6 of the top 20 anti-viral (sensitization) hits were in common, including HIRA and CABIN1, both members of an H3.3 specific chaperone complex.

The additional, recently published genome-wide screens for SARS-CoV-2 host factors have varied in the viral isolate, the CRISPR library, and the cell type (**Table 1**) (Baggen et al., 2021; Daniloski et al., 2021; Schneider et al., 2021; Wang et al., 2021; Wei et al., 2021; Zhu et al., 2021). We acquired the read counts from all these screens and re-processed the data via the same analysis pipeline to enable fair comparisons (see Methods); top-scoring genes were consistent with the analyses provided in the original publications. Two screens, using different CRISPR libraries, were conducted in A549 cells engineered to express ACE2; comparison of these results showed a greater number of statistically significant hits in the Zhang-Brunello dataset (Zhu et al., 2021) compared to the Sanjana-GeCKO dataset (Daniloski et al., 2021), but results were generally consistent between the two, with 10 genes shared in the top 20 (**Figure S1C**). Likewise, three groups conducted survival screens in related cell systems (**Figure S1D)**: Huh7 cells (Daelmans-Brunello (Baggen et al., 2021)); Huh7.5 cells (Poirier-Brunello (Schneider et al., 2021)), a derivative of Huh7, which have biallelic loss-of-function mutation in the DDX58/RIG-I sensor; and Huh7.5.1 cells, engineered to overexpress ACE2 and TMPRSS2 (Puschnik-GeCKO (Wang et al., 2021)). All three screens identified TMEM106B as a top hit, and we observed the best pair-wise correlation between the two screens that used Huh7.5 cells (**Figure S1D**).

**Table 1.** **Properties of SARS-CoV-2 host factor screens assayed by cell viability.** For each library, the number of unique guides per gene is indicated in parentheses. The essential gene QC serves as a metric for screen quality (see Methods) in the untreated arm, when applicable; the number of days post-library introduction until the end of the experiment is written after the semicolon.

| Study | Cell line | Library<br>(#guides/gene) | Viral isolate | Essential gene<br>QC (ROC-AUC;<br>#days) |
| --- | --- | --- | --- | --- |
| Wei,<br>Wilén | Vero E6 | C. sabeus (4) | USA-WA1/2020 | 0.82; 16-18 days |
| This study | Vero E6 | C. sabeus (4) | BetaCoV/France/<br>IDF0372/2020 | 0.84; 20-24 days |
| Daniloski,<br>Sanjana | A549-ACE2 | GeCKO (6) | USA-WA1/2020 | 0.62; 18 days |
| Zhu,<br>Zhang | A549-ACE2 | Brunello (4) | nCoV-SH01 | 0.68; 14+ days |
| Baggen,<br>Daelemans | Huh7 | Brunello (4) | Belgium/GHB-03021/2020 | 0.78; 44 days |
| Schneider, Poirier | Huh7.5 | Brunello (4) | USA-WA1/2020 | 0.92; 12-21 days |
| Wang, Puschnik | Huh7.5.1-ACE2-TMPRSS2 | GeCKO (6) | USA-WA1/2020 | 0.56; 19 days |
| This study | Caco-2-ACE2 | Brunello (4) | BetaCoV/France/<br>IDF0372/2020 | 0.71; 19 days |
| This study | Calu-3 | Gattinara (2) | BetaCoV/France/<br>IDF0372/2020 | 0.84; 13-30 days |
| This study | Calu-3 | Calabrese (6) | BetaCoV/France/<br>IDF0372/2020 | n/a; 17-19 days |

We next averaged gene-level Z-scores and compared results across the Vero E6, A549, and Huh7.5 cell lines. Examining the top 20 genes from each cell line, and using a lenient Z-score threshold of 3 to consider a gene a hit, we generated a Venn diagram to examine their overlap (**Figure 1D**). By these criteria, only ACE2 and CTSL scored in all three models, and 3 additional genes overlapped in two cell lines. Examining the cell-line specific hits, in Vero-E6 cells we continued to observe an enrichment of BAF proteins SMARCA4 and DPF2 (Wei et al., 2021); notably, another nBAF complex member, ARID1A, also scored in A549 cells. Genes scoring uniquely in A549 cells included several COMM domain-containing proteins, which have been implicated in NF-kB signaling (Szklarczyk et al., 2021). Finally, Huh7.5 cells showed specificity for EXT1 and EXT3L, genes involved in heparin sulfate biosynthesis, as well as SLC35B2, which transports PAP, a substrate for intracellular sulfation. Overall, these analyses suggest that individual cell models are particularly suited, in as yet unpredictable ways, to probe different aspects of SARS-CoV-2 host factor biology.

### Whole-genome knockout and activation screens to identify genes regulating SARS-CoV-2 replication in Calu-3 cells

Calu-3 cells, a lung adenocarcinoma cell line, are a particularly attractive model for exploring SARS-CoV-2 biology, as they naturally express ACE2 and TMPRSS2. Furthermore, we have previously shown that Calu-3 cells behave highly similarly to primary human airway epithelia when challenged with SARS-CoV-2 (Rebendenne et al., 2021). Additionally, they are suited to viability-based screens, as they are highly permissive to SARS-CoV-2 and show high levels of cytopathic effects upon replication, although the slow doubling time of the cells (∼5-6 days) presents challenges for scale-up.

To conduct genome-wide CRISPR knockout and activation screens (**Figure 2A**), Calu-3 cells were stably engineered to express Cas9 or dCas9-VP64, respectively. Calu-3-Cas9 cells showed >94% Cas9 activity (**Figure S2A**) and Calu-3-dCas9-VP64 cells transduced to express sgRNAs targeting the *MX1* and *IFITM3* promoters induced expression to a similar magnitude as following interferon-treatment (**Figures S2B-C**). The more compact Gattinara library (DeWeirdt et al., 2020) was selected for the knockout screen due to the difficulty of scaling-up this cell line, while the Calabrese library was used for the CRISPR activation (CRISPRa) screen (Sanson et al., 2018). Cells were transduced with the libraries in biological triplicates at a low MOI, selected with puromycin, and 15 to 18 days post-transduction, were either harvested for subsequent genomic DNA extraction or challenged with SARS-CoV-2 at MOI 0.005, which led to >90% cell death in 3-5 days. The surviving cells were then cultured in conditioned media, expanded and harvested when cell numbers were sufficient for genomic DNA extraction (see Methods). The screening samples were processed and analyzed as above.

**Figure 2.**
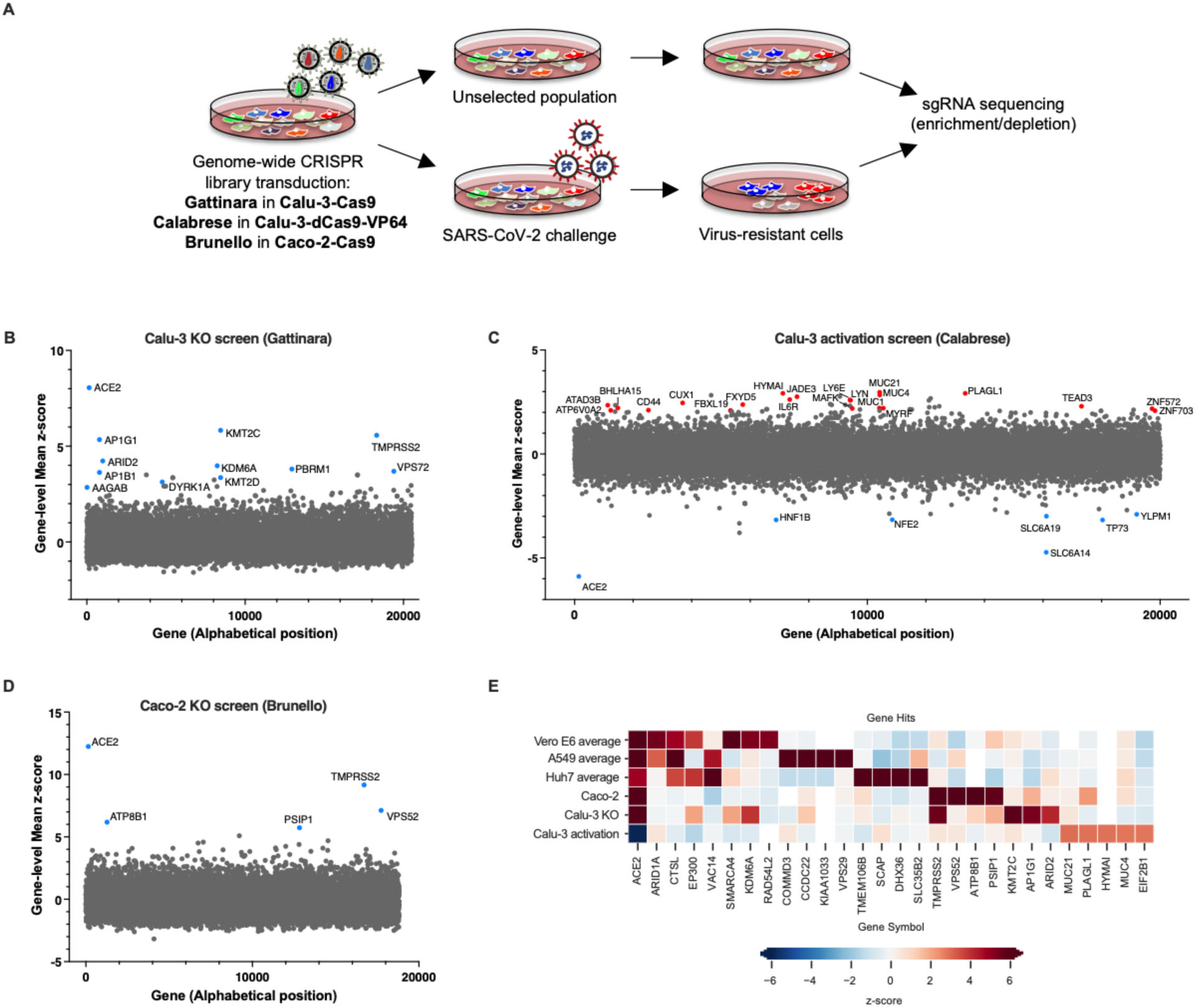
**Genome-wide CRISPR screens in Calu-3 reveal new regulators of SARS-CoV-2.** **A.** Schematic of pooled screens (Calu-3 KO/CRISPRa, Caco-2 KO). **B.** Scatter plot showing the gene-level mean z-scores of genes when knocked out in Calu-3 cells. The top genes conferring resistance to SARS-CoV-2 are annotated and shown in blue. This screen did not have any sensitization hits. **C.** Scatter plot showing the gene-level mean z-scores of genes when over-expressed in Calu-3 cells. The top genes conferring resistance to SARS-CoV-2 are annotated and shown in blue. The top genes conferring sensitivity to SARS-CoV-2 are annotated and shown in red. **D.** Scatter plot showing the gene-level mean z-scores of genes when knocked out in Caco-2 cells. The top genes conferring resistance to SARS-CoV-2 are annotated and shown in blue. **E.** Heatmap of top 5 resistance hits from each cell line after averaging across screens in addition to genes that scored in multiple cell lines based on the criteria used to construct the Venn diagram in Figure 1D (based on (Baggen et al., 2021; Daniloski et al., 2021; Schneider et al., 2021; Wang et al., 2021; Wei et al., 2021; Zhu et al., 2021) and this study).

The knockout screen was most powered to identify proviral factors (**Figure S2D**), and the top three genes were ACE2, KMT2C and TMPRSS2 (**Figure 2B**). Importantly, the latter did not score in any of the cell models discussed above; conversely, CTSL did not score in this screen. Interestingly, whereas the BAF-specific ARID1A scored in Vero E6 cells and A549 cells, PBAF-specific components ARID2 (rank 5) and PRBM1 (rank 7) scored as top hits in Calu-3. Additional new hits include AP1G1 (rank 4), AP1B1 (rank 9), and AAGAB (rank 22), which code for proteins involved in the formation of clathrin-coated pits and vesicles, and are important for vesicle-mediated, ligand-receptor complex intracellular trafficking.

We next examined the CRISPRa screen (**Figures 2C and S2E)**. In contrast to the knockout screen, here we were able to detect both pro- and anti-viral genes; we speculate this is due to the shorter length of time in culture post-SARS-challenge for the activation screens (2 weeks, compared to 4 in the knockout screens). Assuringly, the top-scoring pro-viral (sensitization) hit was ACE2. Several solute carrier (SLC) transport channels also scored on this side of the screen, including SLC6A19 (rank 8), which is a known partner of ACE2 (Camargo et al., 2020). Furthermore, SLC6A14 (rank 2) has been implicated in cystic fibrosis progression and shown to regulate the attachment of *Pseudomonas* to human bronchial epithelial cells (Di Paola et al., 2017). On the antiviral side of the screen, a top scoring hit was LY6E (rank 10), which is a known restriction factor of SARS-CoV-2 (Pfaender et al., 2020), further validating the ability of this screening technology and cellular model to identify known biology. Additionally, MUC21 (rank 1), MUC4 (rank 4), and MUC1 (rank 26) all scored; Mucins are heavily glycosylated proteins and have a well-established role in host defense against pathogens (Chatterjee et al., 2020; McAuley et al., 2017); moreover, MUC4 has been recently proposed to possess a protective role against SARS-CoV-1 pathogenesis in a mouse model (Plante et al., 2020). Finally, we directly compared the knockout and activation screens conducted in Calu-3 cells (**Figure S2G**). The only gene that scored in both the knockout and activation screen, even using a lenient Z-score threshold of >3, was ACE2 emphasizing that different aspects of biology are revealed by these screening technologies.

To expand the range of cell lines examined further, we also performed a knockout screen with the Brunello library in another cell line naturally permissive to SARS-CoV-2 replication, the colorectal adenocarcinoma Caco-2 cell line. Here, however, the cells were engineered to overexpress ACE2 in order to reach sufficient levels of CPE to enable viability-based screening. Similar to Calu-3 cells, ACE2 and TMPRSS2 were the top resistance hits (**Figures 2D, S2D and S2F**), indicating that Caco-2 and Calu-3 cells, unlike previously used models, rely on TMPRSS2-mediated cell entry, rather than the CTSL-mediated endocytic pathway, which did not score in this cell line (Z-score=-0.2). Assembling all the proviral genes identified across 5 cell lines, we observed a continuation of the trend that screen results are largely cell line dependent (**Figure 2E**).

### Individual validations via CRISPR KO confirm the identification of new proviral genes, including members of the AP1 complex

First we focused on the proviral genes identified in our KO screens and selected 22 candidates among the top ones identified in the screens performed in Calu-3, Vero E6 and Caco-2 cells. We designed 2 sgRNAs to target these genes and generated polyclonal knockout Calu-3 cell populations. In parallel, we generated 2 negative control cell lines (coding non-targeting sgRNAs) and 2 positive control cell lines (*ACE2* and *TMPRSS2* KO). Two weeks post-transduction, knockout cell lines were challenged with SARS-CoV- 2 bearing the mNeonGreen reporter (Xie et al., 2020a) and the percentage of infected cells was scored by flow cytometry (**Figure 3A**). The knockout of about half of the selected genes induced at least a 50% decrease in infection efficiency. Among them, *AP1G1* KO had an inhibitory effect as drastic as *ACE2* KO (>95% decrease in infection efficiency), showing an absolutely essential role of this particular gene. Another member of the Adaptin family, *AP1B1,* and a known partner of the AP1 complex, *AAGAB*, also had an important impact, albeit not as strong (∼70-90% decrease in infection). The KO of 3 other genes *KMT2C*, *EP300*, and *ATP8B1*, which code for a lysine methyltransferase, a histone acetyl transferase and a flippase, respectively, inhibited the infection efficiency by at least 50%. In parallel, we tested the impact of candidate knockout on SARS-CoV-2-induced CPEs. Cells were infected with wild-type SARS-CoV-2 at an MOI of 0.005 and colored with crystal violet when massive CPE was observed in the negative controls, ∼5 days post-infection (**Figure S3A**). CPE analyses globally mirrored data obtained with mNG reporter viruses, showing that the identified genes were *bona fide* proviral factors and not genes the KO of which would only protect cells from virus-induced cell death. Encouragingly, based on a recent scRNA-seq study (Chua et al., 2020), the best-validated candidate genes, i.e. *AP1G1*, *AB1B1*, *AAGAB*, *KMT2C*, *EP300* and *ATP8B1*, were all well expressed in SARS-CoV-2 target cells from the respiratory epithelia (**Figure S3B**).

**Figure 3.**
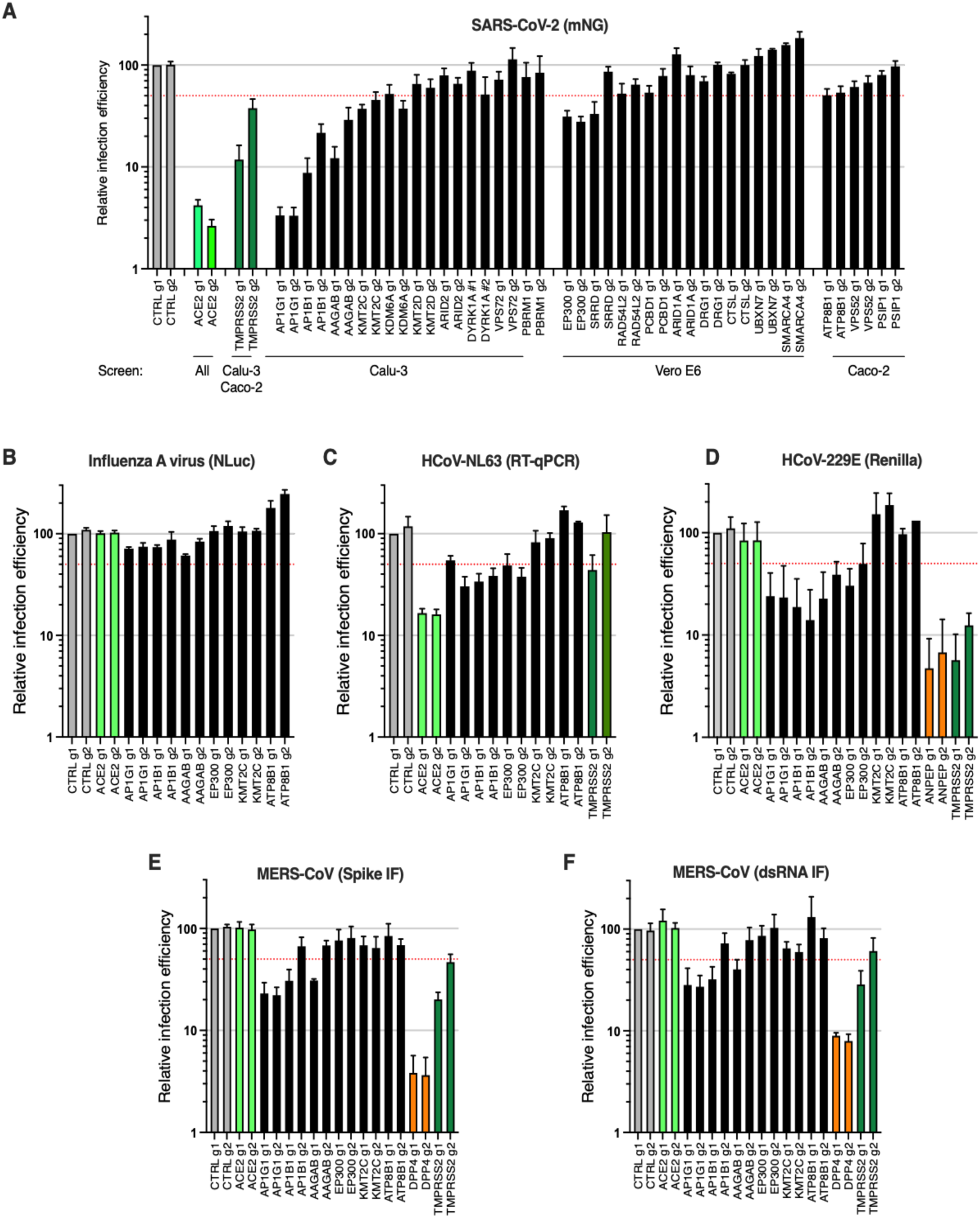
**Impact of the identified proviral genes on coronaviruses SARS-CoV-2, HCoV-229E HCoV-NL63, and MERS-CoV and on orthomyxovirus influenza A.** Calu-3-Cas9 cells were stably transduced to express 2 different sgRNAs (g1, g2) per indicated gene and selected for 10-15 days. **A.** Cells were infected with SARS-CoV-2 bearing the mNeonGreen (mNG) reporter and the infection efficiency was scored 48h later by flow cytometry. The cell line/screen in which the candidates were identified is indicated below the graph. **B.** Cells were infected with influenza A virus bearing the Nanoluciferase (NLuc) reporter and 10h later relative infection efficiency was measured by monitoring Nluc activity. **C.** Cells were infected with HCoV-NL63 and 5 days later, relative infection efficiency was determined using RT-qPCR. **D.** Cells were infected with HCoV-229E-Renilla and 48-72h later, relative infection efficiency was measured by monitoring Renilla activity. **E-F.** Cells were infected with MERS-CoV and 16h later, the percentage of infected cells was determined using anti-Spike (**E**) or anti-dsRNA (**F**) immunofluorescence (IF) staining followed by microscopy analysis (10 fields per condition). The mean and SEM of 4 to 7 independent experiments (A; with the notable exception of the genes with no validated impact in Calu-3 cells, i.e. *DYRK1A*, *VPS72*, *PBRM1*, *DRG1*, *UBXN7*, *CRSL1*, *SMARCA4*, n=2), 4 (B), 3 (D, E, F) or 2 (C) independent experiments. The red dashed line represents 50% inhibition.

We then investigated the effect of these genes on other respiratory viruses. Noteworthy, knockout had no substantial impact on the replication of a respiratory virus from another family, the orthomyxovirus influenza A virus (IAV) strain A/Victoria/3/75 (H3N2) (**Figure 3B**). In contrast, HCoV-NL63 replication was impacted by *AP1G1*, *AP1B1* and *EP300* KO, but not by *KMT2C* or *ATP8B1* KO (**Figure 3C**). Interestingly, seasonal HCoV-229E and highly pathogenic MERS-CoV, which do not use ACE2 for viral entry but ANPEP and DPP4, respectively, were also both strongly affected by *AP1G1*, and, to some extent, by *AP1B1* and *AAGAB* KO (**Figure 3D-F**), showing a pan-coronavirus role of these genes.

Next, we aimed to determine the life cycle step affected by the candidate KOs and we examined the impact of the best validated candidate KO (i.e. with an effect >50% decrease in mNG reporter expression, **Figure 4A**) on ACE2 global expression levels (**Figure 4B**). Immunoblot analysis revealed similar or higher expression levels of ACE2 in the different KO cell lines in comparison to controls, with the exception of *ACE2 and EP300* KO cells, which had decreased levels of ACE2. We then took advantage of recombinant Spike Receptor Binding Domain (RBD) fused to a mouse Fc fragment, in order to stain ACE2 at the cell surface (**Figure 4C**). Using this system, we did not observe a substantial decrease in ACE2 at the plasma membrane, apart from *ACE2 and EP300* KO cell lines, as expected.

**Figure 4.**
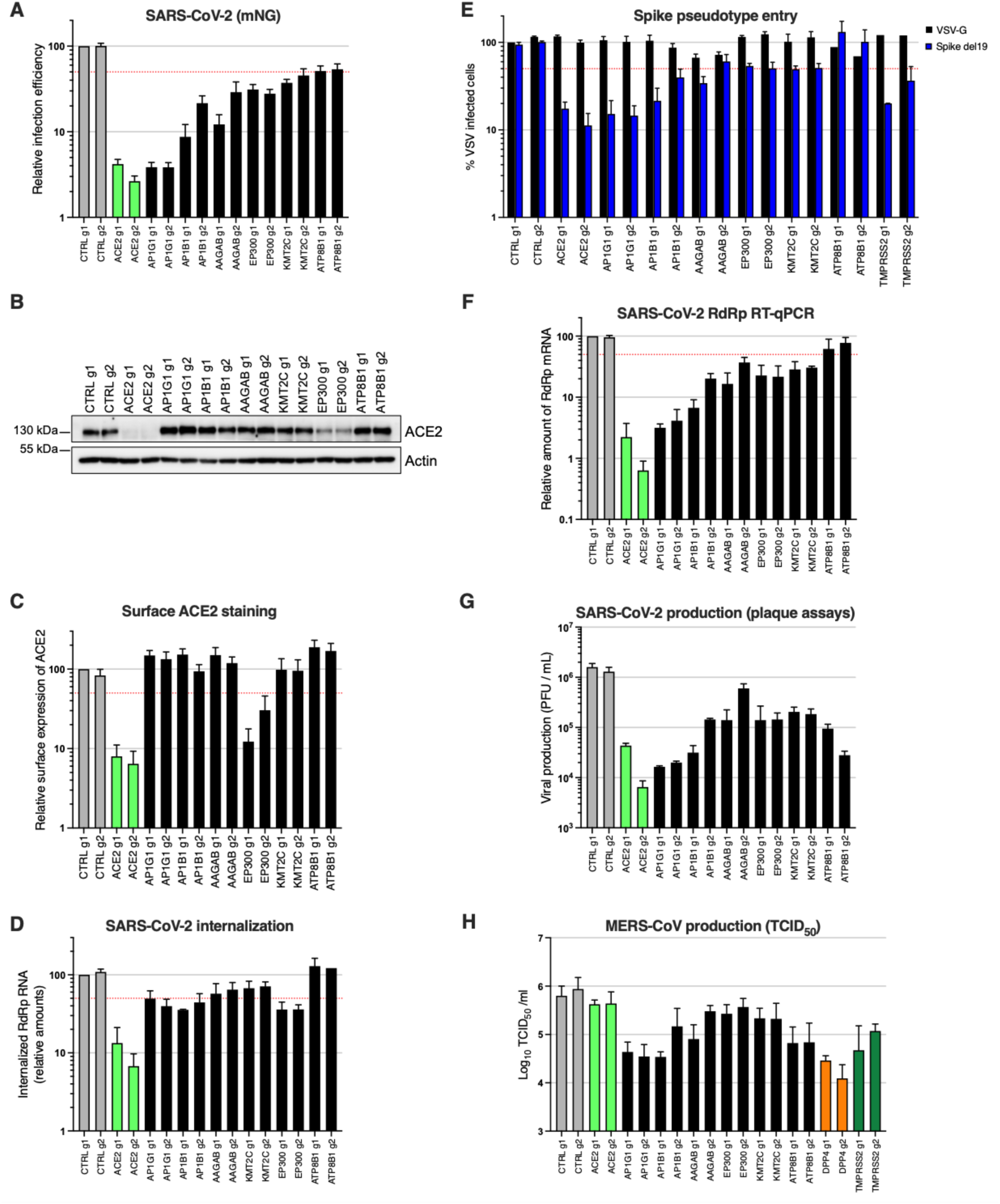
**Characterization of the impact of identified SARS-CoV-2 dependency factors.** Calu-3-Cas9 cells were stably transduced to express 2 different sgRNAs (g1, g2) per indicated gene and selected for 10-15 days. **A.** Cells were infected with SARS-CoV-2 bearing the mNeonGreen (mNG) reporter and the infection efficiency was scored 48h later by flow cytometry. **B.** The expression levels of ACE2 were analyzed by immunoblot, Actin served as a loading control. **C.** Relative surface ACE2 expression was measured using a Spike-RBD-Fc fusion and a fluorescent secondary antibody followed by flow cytometry analysis. **D.** Cells were incubated with SARS-CoV-2 at MOI 5 for 2h at 37°C and then treated with Subtilisin A followed by RNA extraction and RdRp RT-qPCR analysis as a measure of viral internalization. **E.** Cells were infected with Spike del19 and VSV-G pseudotyped, GFP expressing VSV and infection efficiency was analyzed 24h later by flow cytometry. **F.** Cells were infected with SARS-CoV-2 at MOI 0.05 and, 24h later, lysed for RNA extraction and RdRp RT-qPCR analysis. **G.** Aliquots of the supernatants from F were harvested and plaque assays were performed to evaluate the production of infectious viruses in the different conditions. **H.** Cells were infected with MERS-CoV and 16h later, infectious particle production in the supernatant was measured by TCID_50_. The mean and SEM of at least 5 (A), 3 (C, D, E, F H) independent experiments or representative experiments (B and G) are shown. The red dashed line represents 50% inhibition (A, C-F).

In order to assess the internalization efficiency of viral particles, we then incubated the KO cell lines with SARS-CoV-2 at an MOI of 5 for 2h at 37°C, and treated the cells with Subtilisin A in order to eliminate the cell surface-bound viruses, followed by RNA extraction and RdRp RT-qPCR to measure the relative amounts of internalized viruses (**Figure 4D**). This approach showed that *AP1G1*, *AP1B1*, *AAGAB* and *EP300* impacted SARS-CoV-2 internalization to at least some extent, but not *ATP8B1*. We then used VSV particles pseudotyped with SARS-CoV-2 Spike, bearing a C-terminal deletion of 19 aminoacids (hereafter named Spike del19) as a surrogate for viral entry (Schmidt et al., 2020), in comparison to VSV-G pseudotypes (**Figure 4E**). Of note, both *ACE2* and *TMPRSS2* knockout specifically impacted Spike del19-VSV infection, confirming that the pseudotypes mimicked wild-type SARS-CoV-2 entry in Calu-3 cells. We observed that Spike del19-dependent entry was affected in most cell lines in comparison to VSV-G-mediated entry, with, again, the notable exception of *ATP8B1* KO cells, implying a later role for this gene. Analysis of SARS-CoV-2 RNA replication by RdRp RT-qPCR (**Figure 4F**) and viral production in the cell supernatants by plaque assays (**Figure 4G**) perfectly mirrored the data obtained using the mNG virus reporter, apart from *ATP8B1* KO cells. Indeed, in the latter, there was only around 50% decrease in viral RNA replication or mNG reporter expression, but more than one order of magnitude decrease in viral production, suggesting a late block during viral replication (**Figures 4F-G**). Importantly, highly similar results were obtained with MERS-CoV for *AP1G1* and *AP1B1*, which had an impact comparable to *DPP4* receptor KO on viral production (**Figure 4H**). Moreover, as observed for SARS-CoV-2, *ATP8B1* KO also strongly impacted infectious MERS-CoV particle production/release, whereas it had only a minor impact on infection as measured by Spike or dsRNA intracellular staining (**Figures 4H and 3E-F**), arguing for a common and late role of this gene in the coronavirus replicative cycle.

### CRISPR activation screen reveals new anti-SARS-CoV-2 genes, including *Mucins*, *CD44* and *IL6R*

Next, 21 genes among the top-ranking hits conferring resistance to SARS-CoV-2 replication from the CRISPR activation screens were selected for individual validation, using two different sgRNAs in Calu-3-dCas9-VP64 cells. In parallel, non-targeting sgRNAs and sgRNAs targeting *ACE2* and *IFNL2* promoters were used as controls. 10-15 days post-transduction, the sgRNA-expressing cell lines were challenged with SARS-CoV-2 bearing the mNeonGreen reporter, as previously, and the percentage of infected cells was scored by flow cytometry (**Figure 5A**). As expected (Pfaender et al., 2020; Rebendenne et al., 2021; Stanifer et al., 2020), the induction of *IFNL2* and *LY6E* expression potently decreased SARS-CoV-2 replication. We observed that the increased expression of the vast majority of the selected hits induced at least a 50% decrease in infection efficiency with at least 1 of the 2 sgRNAs. Some genes had a particularly potent impact on SARS-CoV-2 and decreased the replication levels by 80-90% or more, including the Mucin genes *MUC1*, *MUC21*, *MUC4*, as well as *CD44*, *PLAGL1*, *IL6R*, *TEAD3* and *LYN* (**Figure 5A**). *CD44* codes for a cell surface transmembrane glycoprotein playing multiple roles in adhesion, cell proliferation and survival, signaling, migration, or lymphocyte activation (Jordan et al., 2015; Ponta et al., 2003), and is particularly well expressed in secretory cells from the airway epithelia (Chua et al., 2020). *PLAGL1* codes for a zinc finger transcription factor that promotes cell cycle arrest and apoptosis through multiple pathways (Vega-Benedetti et al., 2017). *IL6R* (also known as gp80 or CD126) codes for a membrane-bound as well as a soluble receptor for IL6; IL6R is bound to gp130 (or CD130), which mediates signal transduction. Upon binding of IL6 to IL6R, the homodimerization of gp130 is induced and a hexameric complex constituted of IL6/IL6R/gp130 is formed, which induce a signaling cascade through the JAK/STAT and SHP-2/ERK MAPK pathways regulating a variety of biological activities, including host defense (Mihara et al., 2011). *TEAD3* codes for a member of transcriptional enhancer factor (TEF) family of transcription factors and plays roles in development, cell differentiation as well as proliferation (Han et al., 2020; Imajo et al., 2015). *LYN* codes for a membrane-anchored src tyrosine kinase, localized on the cytoplasmic side of the plasma membrane and is an important regulator of signal transduction (Brodie et al., 2018). Noteworthy, LYN was shown to regulate inflammatory responses to bacterial infection (Li et al., 2014) and to be important for flavivirus egress (Li et al., 2020). Additionally, an scRNA seq study had shown that most of the antiviral genes identified here were expressed in a substantial percentage of epithelial cells from the respiratory epithelium, including ciliated cells and secretory cells, the main targets of SARS-CoV-2 (**Figure S4**, based on (Chua et al., 2020)).

**Figure 5.**
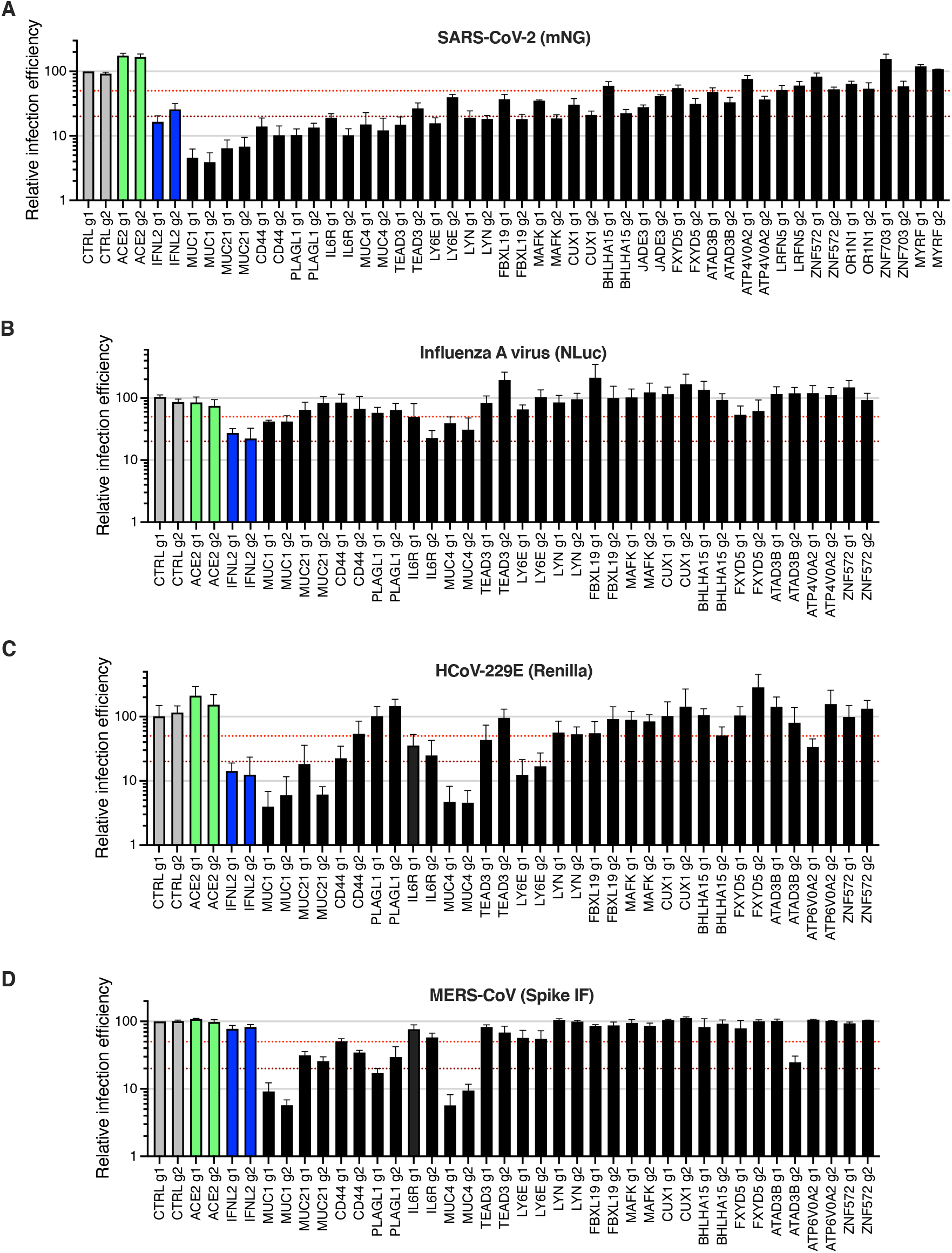
**Impact of the identified antiviral genes on coronaviruses SARS-CoV-2, HCoV-229E, and MERS-CoV, and on orthomyxovirus influenza A.** Calu-3-dCas9-VP64 cells were stably transduced to express 2 different sgRNAs (g1, g2) per indicated gene promoter or negative controls (CTRL) and selected for at least 10-15 days. **A.** Cells were infected with SARS-CoV-2 bearing the mNeonGreen (mNG) reporter and the infection efficiency was scored 48 h later by flow cytometry. **B.** Cells were infected with influenza A virus bearing the Nanoluciferase (NLuc) reporter and 10h later relative infection efficiency was measured by monitoring Nluc activity. **C.** Cells were infected with HCoV-229E-Renilla and 48-72h later, relative infection efficiency was measured by monitoring Renilla activity. **D.** Cells were infected with MERS-CoV and 16h later, the percentage of infected cells was determined using anti-Spike IF staining followed by microscopy analysis (10 fields per condition). The mean and SEM of at least 4 (A) or 3 (B, C, D) independent experiments are shown. The red and dark red dashed lines indicate 50% and 80% inhibition, respectively.

Looking at the antiviral breadth of the validated genes, we observed that the induction of most of them had no impact on IAV infection (**Figure 5B**), with the exception of *MUC4* and *MUC1,* which decreased the infection efficiency by ∼60-70%, as seen previously (McAuley et al., 2017), and *IL6R,* with one of the 2 sgRNAs leading to 75% decrease in infection efficiency. Interestingly, similarly to SARS-CoV-2, HCoV-229E appeared highly sensitive to the increased expression of *MUC*s, *IL6R*, *LY6E*, but was less affected or not affected at all by the other genes, such as *CD44* or *PLAGL1* (**Figure 5C**). MERS-CoV infection was impacted by the 3 Mucin genes of interest and to some extent by *PLAGL1*, *CD44*, *IL6R*, *LY6E* and *ATAD3B*, but not by the other candidates (**Figure 5D**).

Next, we tested the impact on SARS-CoV-2 of some of the best candidates in naturally permissive Caco-2 cells and in A549 cells engineered to ectopically express ACE2 (**Figure S5**). We observed that *MUC4*, *MUC1, MUC21* induction potently decreased SARS-CoV-2 infection in these two other cell lines. Moreover, *PLAGL1* also had a strong impact in A549-ACE2 cells but not in Caco-2 cells, and the opposite was true for *LYN*.

This might suggest a potential cell type specificity for the former (e.g. lung origin) and possibly a dependence on ACE2/TMPRSS2 endogenous expression for the latter. *CD44* and *LY6E* also had an inhibitory effect to some extent in both cell lines. Taken together, this globally showed that the inhibitory effect of the validated candidates is not restricted to Calu-3 cells and can be observed in other cell types.

We then explored the life cycle step affected by antiviral gene expression. The SARS-CoV-2 internalization assay, performed as previously, showed that most of the validated genes, including those showing the strongest inhibitory phenotypes (namely *MUC1*, *MUC21*, *CD44*, *PLAGL1*, *IL6R*, *MUC4*, and *LYN*) impacted viral internalization (**Figure 6A**). The measure of viral entry using Spike del19- or G-pseudotyped VSV particles globally mirrored the internalization data, and showed that G-dependent entry was as sensitive as Spike del19-dependent entry to the induced expression of Mucins, IL6R or LYN (**Figure 6B**). However, we observed that whereas *CD44* and *PLAGL1* had an impact on SARS-CoV-2 entry as measured by our internalization assay (as well as a number of other genes such as *TEAD3*, but with milder effects), there was no effect of these genes on Spike del19-VSV pseudotypes, perhaps highlighting subtle differences in the mechanism of entry by the latter compared to wild-type SARS-CoV-2. Moreover, *LY6E* induction had no measurable impact on viral entry, either using the internalization assay or the VSV pseudotype assay, contrary to what was reported before (Pfaender et al., 2020). Differences in the experimental systems used could explain the differences observed here and would require further investigation. Finally, the impact of the best candidates on SARS-CoV-2 and MERS-CoV replication, measured by RdRp RT-qPCR (**Figure 6C**) and plaque assays (**Figure 6D**) for SARS-CoV-2, or TCID_50_ for MERS-CoV (**Figure 6E**), recapitulated what was observed with SARS-CoV-2 mNG reporter (**Figure 5A**) and MERS-CoV Spike intracellular staining (**Figure 5D**).

**Figure 6.**
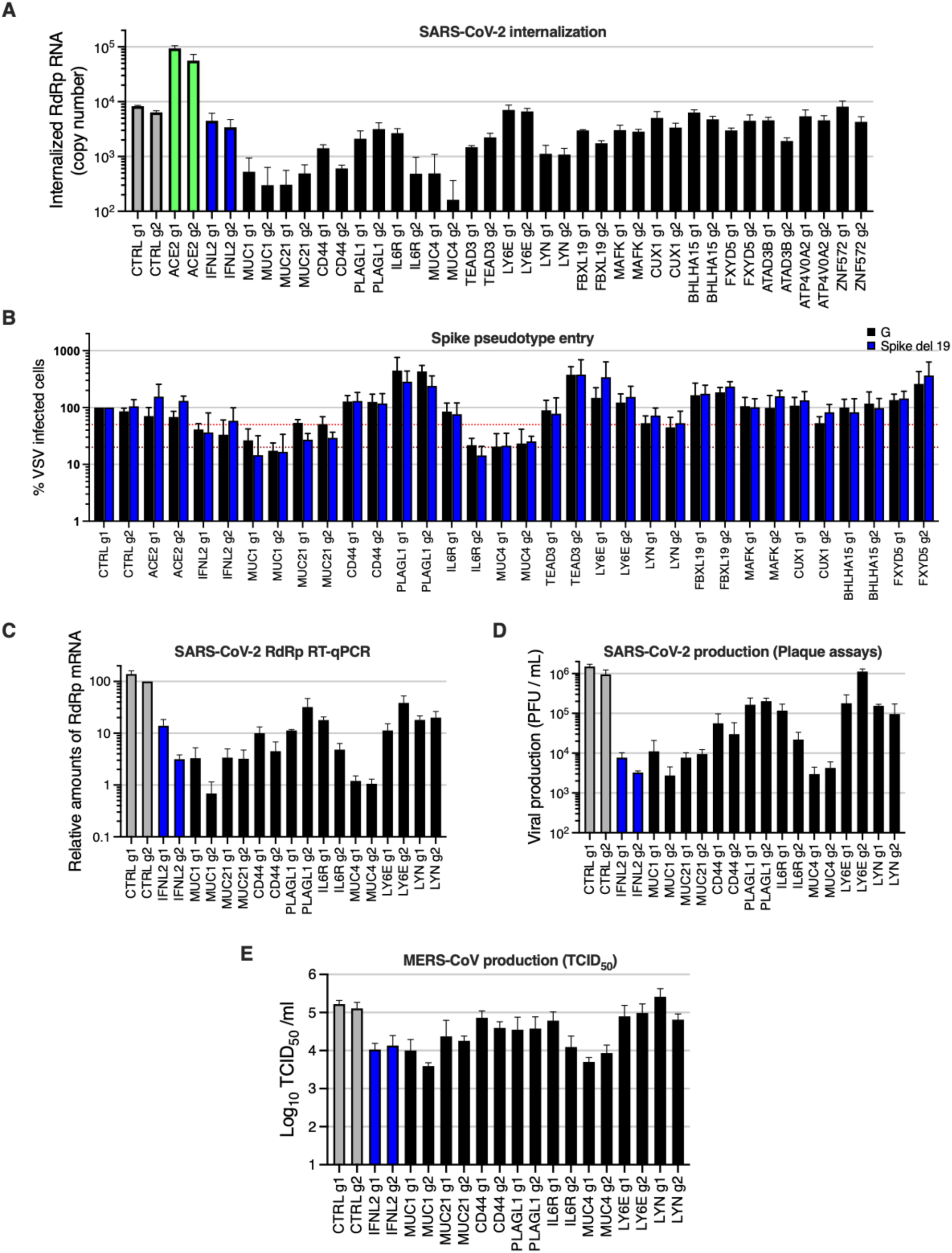
**Characterization of the impact of identified SARS-CoV-2 dependency factors.** Calu-3-dCas9-VP64 cells were stably transduced to express 2 different sgRNAs (g1, g2) per indicated gene promoter and selected for 10-15 days. **A.** Cells were incubated with SARS-CoV-2 at MOI 5 for 2h and then treated with Subtilisin A followed by RNA extraction and RdRp RT-qPCR analysis. **B.** Cells were infected with Spike del19 and VSV-G pseudotyped, Firefly-expressing VSV and infection efficiency was analyzed 24h later by monitoring Firefly activity. The red and dark red dashed lines represent 50% and 80% inhibition, respectively. **C.** Cells were infected with SARS-CoV-2 at MOI 0.05 and, 24h later, lysed for RNA extraction and RdRp RT-qPCR analysis. **D.** Aliquots of the supernatants from C were harvested and plaque assays were performed to evaluate the production of infectious viruses in the different conditions. A representative experiment is shown. **E.** Cells were infected with MERS-CoV and 16h later, infectious particle production in the supernatant was measured by TCID_50_. The mean and SEM of 3 (A, C), 4 (B), 2 (D, E) independent experiments are shown.

Noteworthy, the 3 Mucins of interest had the strongest impact on both SARS-CoV-2 and MERS-CoV production (∼2 log and ∼1 log decrease, respectively, as compared to the controls). The activation of *IL6R*, *CD44*, *PLAGL1*, and *LYN* also had a substantial impact on SARS-CoV-2 replication (∼1 log decrease or more, for at least 1 out of the 2 sgRNAs) but had a globally milder impact on MERS-CoV replication, with *LYN* having no impact at all (**Figure 6D-E**). Whereas Mucins are well-known to act as antimicrobial barriers (Dhar and McAuley, 2019; Linden et al., 2008), the role of the other potent antiviral genes, such as *IL6R*, *CD44* or *PLAGL1*, in limiting SARS-CoV-2 entry remains to be elucidated.

### Individual validations via CRISPR activation reveal additional pro-SARS-CoV-2 genes, including *TP73* and *NFE2*

In addition to the proviral genes identified by the KO screens, we selected several of the top-ranking hits conferring sensitization to SARS-CoV-2 replication in the CRISPR activation screens. Solute carriers SLC6A14 and SLC6A19, transcription factors Tumor Protein P73 (TP73), Hepatocyte nuclear factor-1β (HNF1B) and Nuclear Factor, Erythroid 2 (NFE2) were chosen for individual validations, using two different sgRNAs in Calu-3- dCas9-VP64 cells in parallel to controls, as previously. At least 12-15 days post-transduction, the sgRNA-expressing cell lines were challenged with SARS-CoV-2 bearing a NLuc reporter (Xie et al., 2020b) and the relative infection efficiency was analyzed by monitoring NLuc activity (**Figure 7A**). Among the tested candidates, *TP73*, *HNF1B*, and *NFE2* had the strongest positive impact on SARS-CoV-2 replication (∼3-4-fold increase), which was comparable to what was observed with ACE2 overexpression. *SLC6A19* induction had a slight positive effect on SARS-CoV-2 infection (∼1.5-2-fold). Surprisingly, the induced-expression of *SLC6A14*, which was the top-ranking sensitizing hit after ACE2, had an inhibitory effect on SARS-CoV-2 infection rather than a positive effect, when measuring NLuc reporter activity. However, SARS-CoV-2-induced CPEs were increased in *SLC6A14*-induced cells compared to the control, suggesting a late impact of this gene on viral replication and/or an increase in cell death (**Figure 7B**). Interestingly, none of the identified proviral factors had a positive impact on influenza A virus infection, with the notable exception of *HNF1B*, which had a slight positive impact (**Figure 7C**). In contrast, all the identified proviral genes had a positive impact on HCoV-NL63 infection (**Figure 7D**). We then studied the impact of the candidates on HCoV-229E, using in parallel 2 sgRNAs targeting *ANPEP* as positive controls (**Figure 7E**). Calu-3 cells are known to express low levels of ANPEP (Funk et al., 2012), and, as expected, *ANPEP* receptor induction greatly increased HCoV-229E infection in Calu-3 cells. Among the genes having a positive impact on SARS-CoV-2 and HCoV-NL63, only *TP73* induction had a positive effect on HCoV-229E infection (**Figure 7E**).

**Figure 7.**
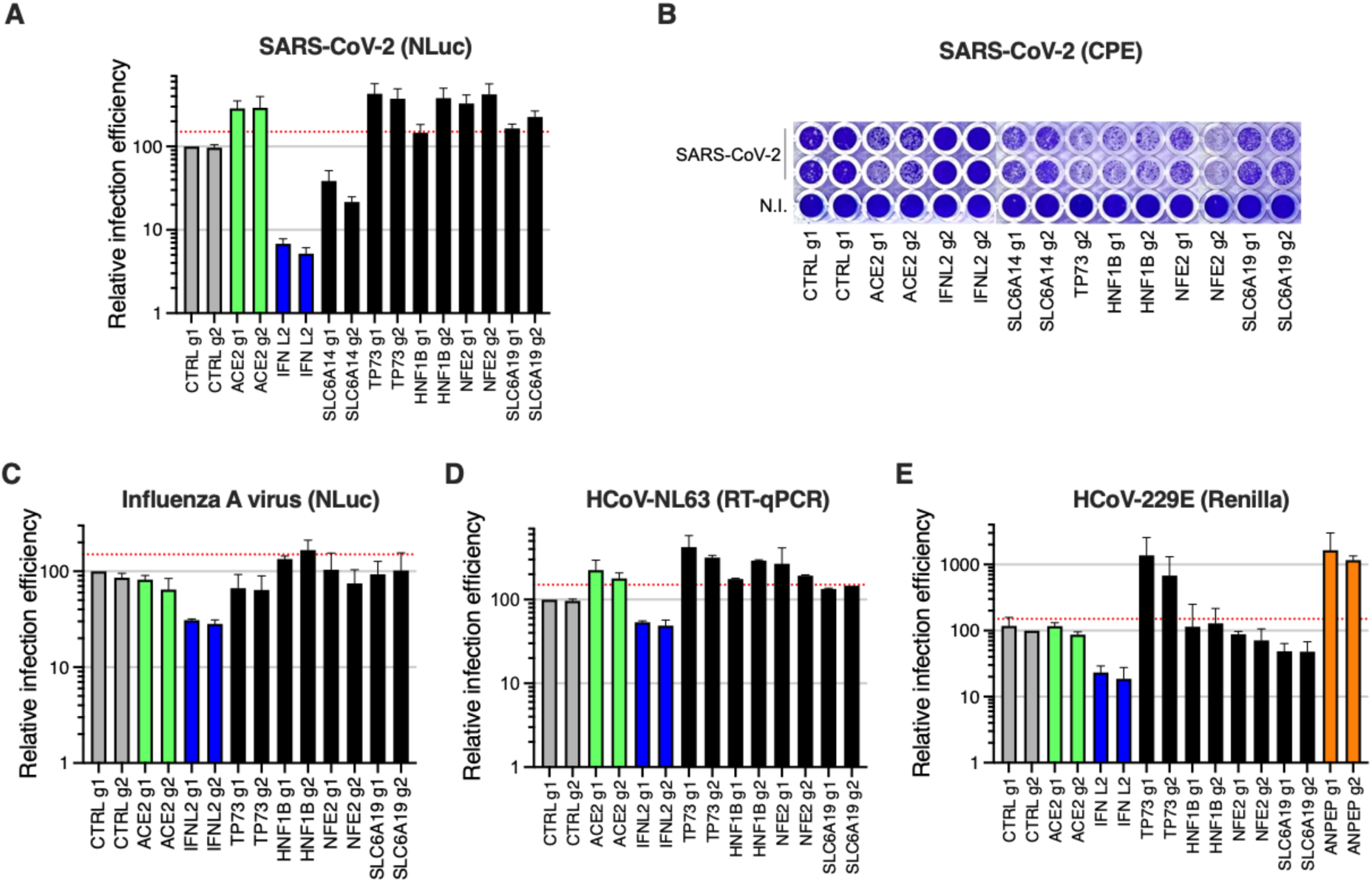
**Impact of the proviral genes identified by CRISPRa on coronaviruses SARS-CoV-2, HCoV-229E, HCoV-NL63 and on orthomyxovirus influenza A.** Calu-3-dCas9-VP64 cells were stably transduced to express 2 different sgRNAs (g1, g2) per indicated gene promoter and selected for 10-15 days. **A.** Cells were non infected (N.I.) or incubated with SARS-CoV-2 bearing NLuc reporter and the infection efficiency was scored 30 h later by monitoring NLuc activity. **B.** Cells were infected by SARS-CoV-2 at MOI 0.05 and ∼5 days later stained with crystal violet. Representative images from 2 independent experiments are shown. **C.** Cells were infected with influenza A virus bearing NLuc reporter and 10h later, relative infection efficiency was measured by monitoring NLuc activity. **D.** Cells were infected with HCoV-NL63 and 5 days later, infection efficiency was determined using RT-qPCR. **E.** Cells were infected with HCoV-229E-Renilla and 72h later, relative infection efficiency was measured by monitoring Renilla activity. The mean and SEM of 4 (A), 3 (C, E) or 2 (D) independent experiments or representative images (B) are shown. The red dashed line indicates 1,5-fold increase in infection efficiency.

In order to decipher the step(s) affected by the induction of the identified proviral genes, we used SARS-CoV-2 internalization and VSV pseudotype assays, as previously (**Figures 8A-B**). Using these 2 assays, we observed that induction of both *HNF1B* and *NFE2* improved viral entry, but not *TP73* or *SLC6A19*, which was surprising for the latter as it is a known partner of ACE2 (Camargo et al., 2020). In line with this, we observed that, despite differences in ACE2 levels in the 2 negative control cell lines, induction of *HNF1B* and *NFE2* seemed to increase ACE2 expression, contrary to that of *TP73* or *SLC6A19* (**Figure S6A**). *TP73* and *SLC6A19* induction, however, increased SARS-CoV-2 RdRp RNA amounts in infected cells as well as infectious particle production, arguing for a post-entry impact on replication (**Figures 8C-D**). Interestingly, the pan-coronavirus cofactor *TP73* (**Figure 7**) was particularly well expressed in ciliated cells from the respiratory epithelium, and SARS-CoV-2 infection in patients positively modulated its expression (**Figure S6B**; (Chua et al., 2020)). TP73 is known to be a pro-apoptotic transcription factor, inducing apoptosis upon DNA damage and regulating DNA damage repair (Stiewe and Pützer, 2001; Urist et al., 2004; Zaika et al., 2011). However, here we show that TP73 does not just play a role in enhancing SARS-CoV-2-induced cell death, as its induction increases viral replication and production. TP73 could be acting indirectly, through the induced expression of SARS-CoV-2 cofactors. Interestingly, although expressed in a lower percentage of cells as compared to *TP73*, *HNF1B* expression was also upregulated in ciliated cells from COVID-19 patients compared to healthy controls (Chua et al., 2020) (**Figure S6B**). HNF1B is a homeodomain containing transcription factor that regulates tissue-specific gene expression positively or negatively, and HNF1B has been shown to modulate lipid metabolism (Long et al., 2017), which might be related to its positive role on SARS-CoV-2 entry, in addition to the observed increase of ACE2 expression. NFE2 is a transcription factor involved in erythroid and megakaryocytic maturation and differentiation and, together with MAFK (which was identified as an antiviral gene by our CRISPRa screen, **Figures 2C** and **5A**), forms a complex, which regulates various pathways (Katsuoka and Yamamoto, 2016). Interestingly, genes regulated by MAFK and NFE2 were both identified as differentially expressed upon SARS-CoV-1 replication (Kumar et al., 2020).

**Figure 8.**
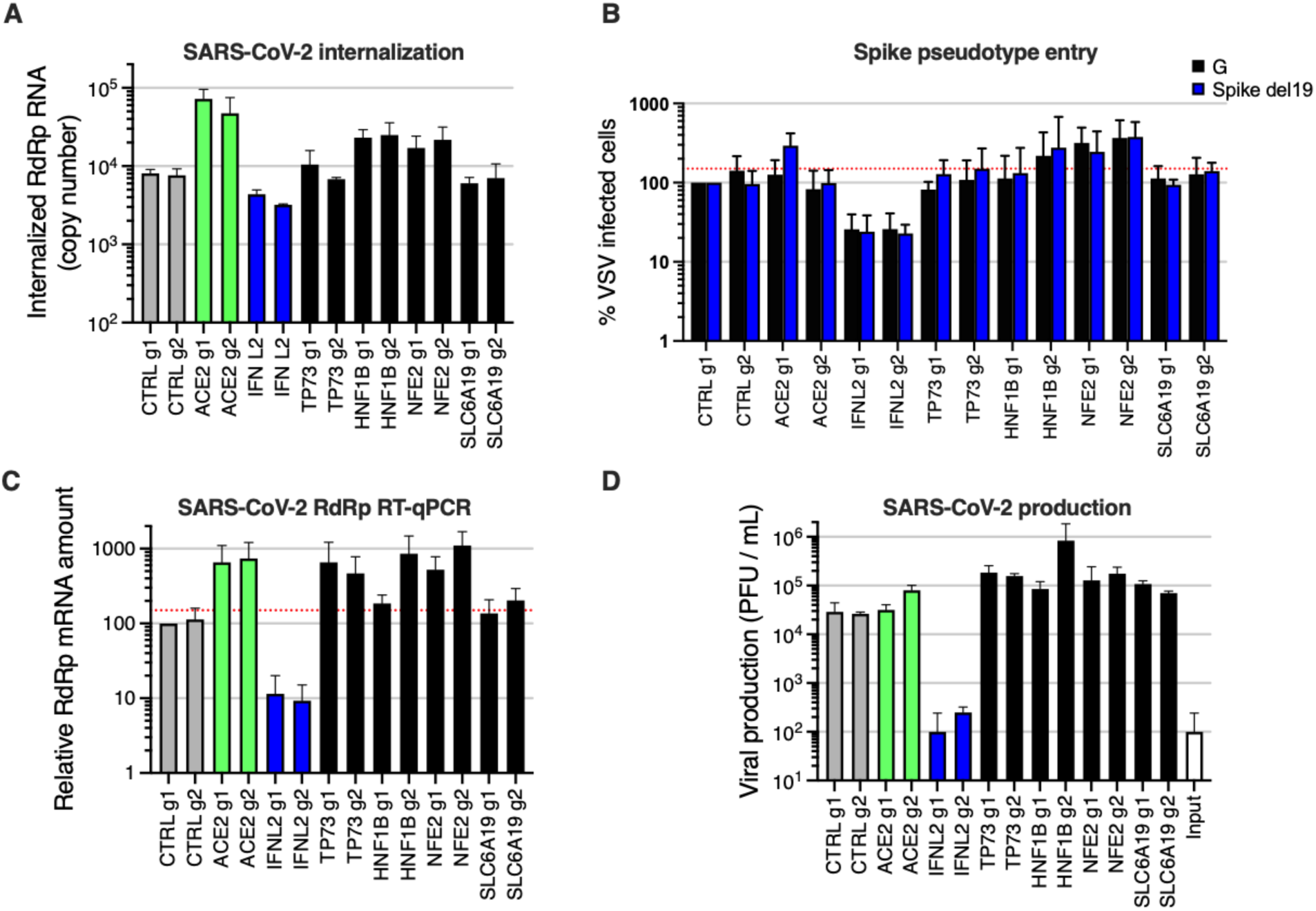
**SARS-CoV-2 life cycle steps affected by the proviral gene induction.** Calu-3-dCas9-VP64 cells were transduced to express 2 different sgRNAs (g1, g2) per indicated gene promoter and selected for 10-15 days. **A.** Cells were incubated with SARS-CoV-2 at MOI 5 for 2h at 37°C and then treated with Subtilisin A followed by RNA extraction and RdRp RT-qPCR analysis as a measure of viral internalization. **B.** Cells were infected with Spike del19 and VSV-G pseudotyped, Firefly-expressing VSV and infection efficiency was analyzed 24h later by monitoring Firefly activity. **C.** Cells were infected with SARS-CoV-2 at MOI 0.05 and, 24h later, lysed for RNA extraction and RdRp RT-qPCR analysis. **D.** Aliquots of the supernatants from C were harvested and plaque assays were performed to evaluate the production of infectious viruses in the different conditions. The mean and SEM of 3 (A, B, C) or 2 (D) independent experiments are shown. The red dashed line indicates 1,5-fold increase in infection efficiency.

## Discussion

Despite intense research efforts, much remains to be discovered about the host factors regulating replication of SARS-CoV-2 and other coronaviruses. Recently, a number of whole-genome CRISPR KO screens successfully identified coronavirus host-dependency factors pandemics (Baggen et al., 2021; Daniloski et al., 2021; Schneider et al., 2021; Wang et al., 2021; Wei et al., 2021; Zhu et al., 2021). However, most of these screens relied on ACE2 ectopic expression and were performed in cells which do not express TMPRSS2, an important cofactor for entry (Hoffmann et al., 2020) (with one notable exception, which relied on TMPRSS2 ectopic expression, (Wang et al., 2021)). A meta-analysis of these screens revealed a high-level of cell type specificity in the hits identified, indicating a need to pursue such efforts in other model cell lines, in order to better define the landscape of SARS-CoV-2 cofactors. In the present study, we performed bidirectional, genome-wide screens in physiologically relevant lung Calu-3 cells, as well as KO screens in intestinal Caco-2 cells. We identified new host-dependency factors, which are not only essential for SARS-CoV-2 replication but also for other coronaviruses, namely MERS-CoV, HCoV-229E and HCoV-NL63. Furthermore, our study unraveled new antiviral genes, some of them with potent and/or broad anti-coronavirus activity.

Simultaneously to our screens, similar bidirectional, genome-wide screens were performed in Calu-3 cells by P. Hsu and colleagues (Biering et al., 2021). Comparisons between our data sets and theirs showed a very good overlap in the hits identified, both in the KO and activation screens (**Figure S7**), with shared hits including host-dependency factors Adaptins AP1G1 and AP1B1 as well as Mucins as antiviral proteins. Interestingly, ATP8B1, which was identified in our Caco-2 KO screen, scored within the 25 best hits in Hsu and colleagues’ Calu-3 KO screen, showing the complementarity of our data. This comparison emphasizes the reproducibility of CRISPR screens conducted across different labs, even when different libraries are used, while further highlighting that the cellular model is the primary source of variability.

Interestingly, we observed that most of the identified genes impacted the early phases of the replicative cycle. This observation was true for both the host dependency factors and the antiviral inhibitors, presumably emphasizing the fact that viral entry is the most critical step of the viral life cycle and probably, as such, the most easily targeted by natural defenses. Among the host-dependency factors essential for viral entry, the Adaptin AP1G1 and, to a lower extent, Adaptin AP1B1 and their partner AAGAB, surprisingly played a crucial role. The AP-1 complex regulates polarized sorting at the trans-Golgi network and/or at the recycling endosomes, and may play an indirect role in apical sorting (Nakatsu et al., 2014). Interestingly, AAGAB has been shown to bind to and stabilize AP1G1, and in *AAGAB* KO cells, AP1G1 is known to be less abundant (Gulbranson et al., 2019), which may suggest a role of AAGAB *via* the regulation of AP-1 complex here. Our data showed that the KO of *AP1G1*, *AP1B1* or *AAGAB* impacted SARS-CoV-2 entry, while not affecting ACE2 expression at the cell surface. In line with this observation, the KO of these factors also impacted MERS-CoV and HCoV-229E, which use different receptors. However, all these coronaviruses use TMPRSS2 for Spike priming in Calu-3 cells, therefore a possible explanation could be that the AP-1 complex might be important for surface expression of TMPRSS2. Alternatively, the AP-1 Adaptins might be important for the proper localization of other plasma membrane components, which play a role in SARS-CoV-2 attachment and/or entry.

Our analysis revealed that another cofactor affecting viral entry, EP300, which is a histone acetyltransferase, was most likely having an indirect effect on SARS-CoV-2 replication, by regulating ACE2 expression. The fact that EP300 impacted HCoV-NL63 but not HCoV-229E or MERS-CoV reinforced this hypothesis. This was also true for two proviral factors identified through our CRISPRa screens, HNF1B and NFE2. In contrast, proviral factor TP73 had no effect on ACE2 expression or viral entry, and actually impacted the 4 coronaviruses we tested here, suggesting the potential regulation of pan-coronavirus factor(s) by this transcription factor.

An exception among the proviral genes that we characterized was ATP8B1, the only one acting at a late stage of the viral life cycle. ATP8B1 belongs to the P4-Type subfamily of ATPases (P4-ATPases) transporters, which are flippases translocating phospholipids from the outer to the inner leaflet of membrane bilayers (Paulusma and Oude Elferink, 2005). ATP8B1 has been shown to be essential for proper apical membrane structure and mutations of this gene have been linked to cholestasis. The fact that ATP8B1 was important for both SARS-CoV-2 and MERS-CoV replication highlighted a potentially conserved role for coronaviruses and it would be of high interest to understand the underlying molecular mechanisms. Interestingly, ATP8B1 and its homologous ATP8B2 were recently identified as binding-partners of SARS-CoV-2 ORF3 and M, respectively (Stukalov et al., 2021), suggesting that the virus might subvert their functions. Of note, TMEM41B, an integral protein of the endoplasmic reticulum known to regulate the formation of autophagosomes, lipid droplets and lipoproteins, was recently shown to be both an essential coronavirus cofactor (Schneider et al., 2021) and a phospholipid scramblase whose deficiency impaired the normal cellular distribution of cholesterol and phosphatidylserine (Li et al., 2021). Whether ATP8B1 depletion could play a similar role in coronavirus replication remains to be determined.

Among the best antivirals we identified through our CRISPR activation screens, the well-known antimicrobial defenses, membrane-associated Mucins played a broad and potent role at limiting coronavirus entry. Interestingly, these Mucins were upregulated in COVID-19 patients (Chua et al., 2020). Additionally, we showed that induced expression of two other membrane proteins, CD44 and IL6R, could also limit SARS-CoV-2 viral entry. Both these proteins are classically seen as important players during immune responses, being involved mainly in adhesion/trafficking and pro-inflammatory processes, respectively. Interestingly, CD44 has also been demonstrated to serve as a platform that brings other membrane receptors together with actin cytoskeleton, possibly within lipid rafts (Jordan et al., 2015). One can thus hypothesize that CD44 might prevent virus entry by acting on specific cellular membrane domains. Regarding IL6R, it is interesting to note that Tocilizumab, a monoclonal antibody against this protein, has been used in clinical trials in severe COVID-19 patients. Indeed, IL-6 is one of the major cytokines responsible for the exacerbated inflammation observed in severe COVID-19 patients. However, Tocilizumab did not improve the clinical outcome of severe COVID-19 (Rosas et al., 2021). Although the exact molecular mechanism of action of how overexpressed IL6R prevents SARS-CoV-2 entry remains to be uncovered, IL6R signaling might indirectly protect lung epithelial cells from infection *via* the induction of innate defenses.

In conclusion, our study unraveled a new network of SARS-CoV-2 and other coronavirus regulators, in model cell lines physiologically expressing ACE2 and TMPRSS2. Importantly, the main natural targets of SARS-CoV-2 in the respiratory tract do co-express ACE2 and TMPRSS2 (Chua et al., 2020; Valyaeva et al., 2020), which highlight the importance of the models used here. Further characterization work on this newly identified landscape of coronavirus regulators might guide future therapeutic intervention.

## Materials and Methods

### Plasmids and constructs

The lentiviral vector expressing ACE2 (pRRL.sin.cPPT.SFFV/ACE2, Addgene 145842) has been described (Rebendenne et al., 2021). The pLX_311-Cas9 (Addgene 96924) and pXPR_BRD109, which express Cas9 and dCas9-VP64, respectively, have been described (Sanson et al., 2018). LentiGuide-Puro vector was a gift from Feng Zhang (Sanjana et al., 2014; Shalem et al., 2014) (Addgene 52963) and we have described before the LentiGuide-Puro-CTRL g1 and g2 (Doyle et al., 2018) (Addgene 139455 and 139456). pXPR_502 vector for sgRNA expression for CRISPRa was also described (Sanson et al., 2018) (Addgene 96923). Guide RNA coding oligonucleotides were annealed and ligated into BsmBI-digested LentiGuide-Puro or pXPR_502 vectors, as described (Addgene). See **Tables 2 and 3** for the sgRNA coding sequences used. pcDNA3.1_spike_del19 was a gift from Raffaele De Francesco (Addgene 155297).

**Table 2.**
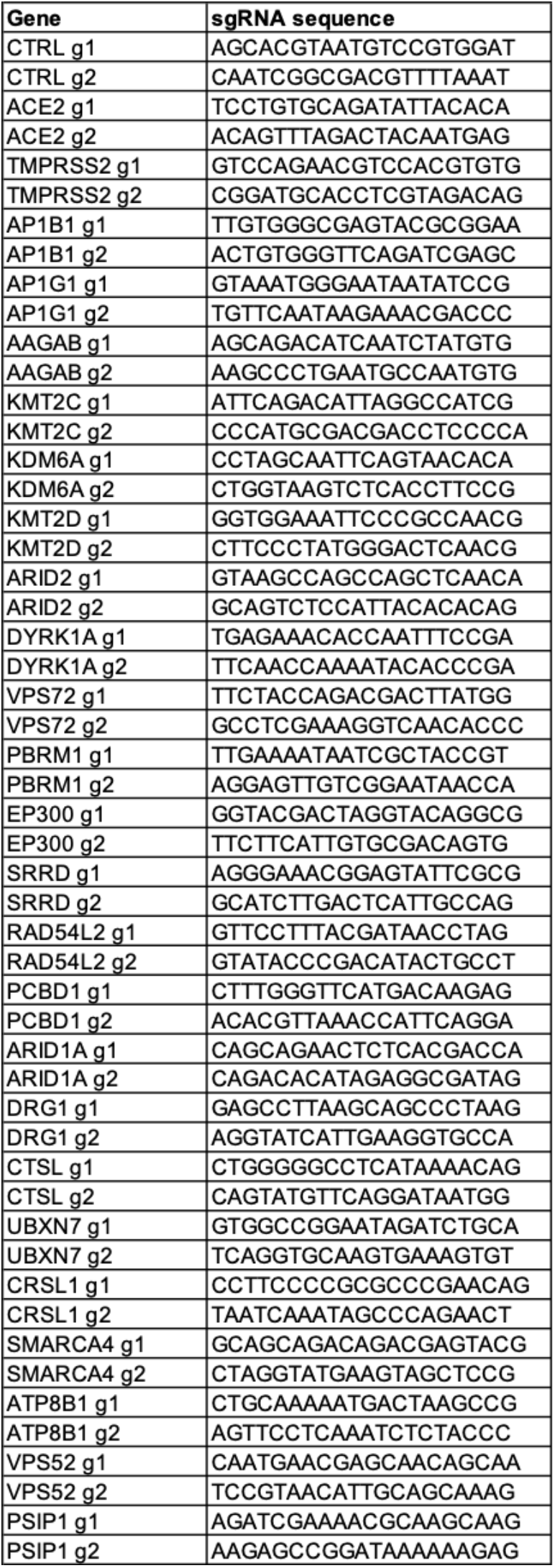
sgRNA sequences used for CRISPR KO perturbations

**Table 3.**
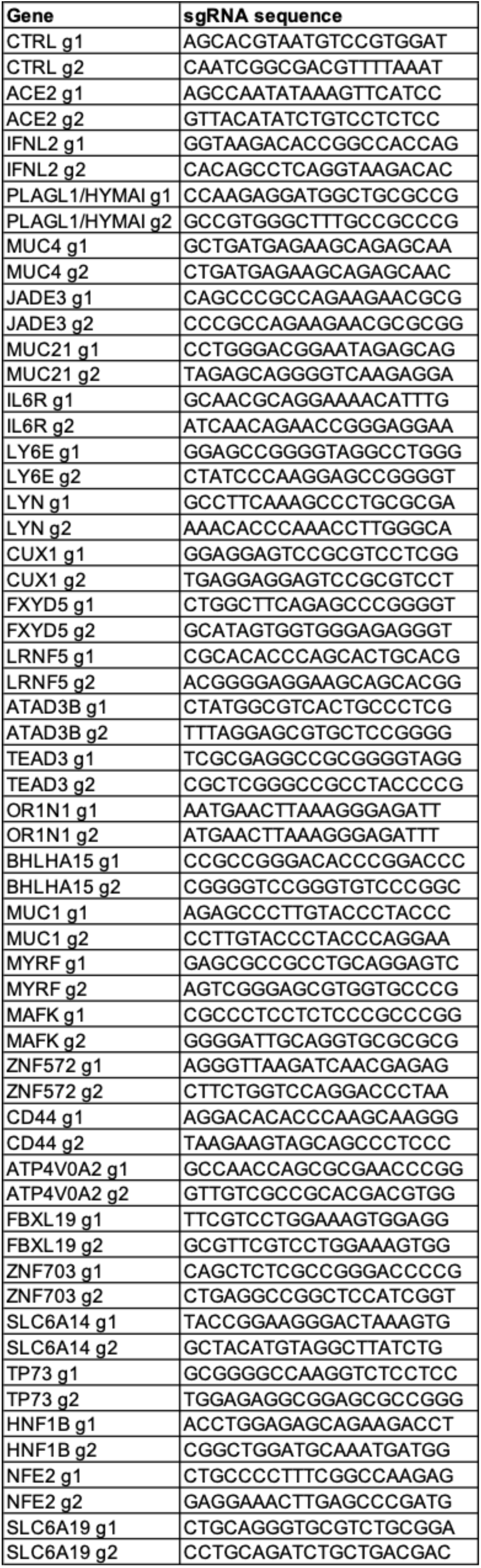
sgRNA sequences used for CRISPRa perturbations

### Cell lines

Human HEK293T, Caco-2, Calu-3, A549, Huh-7, and Huh7.5.1, simian Vero E6 and LLC-MK2, dog MDCK cells were maintained in complete Dulbecco’s modified Eagle medium (DMEM) (Gibco) supplemented with 10% foetal bovine serum and penicillin/streptomycin. Human Caco-2 and Calu-3, simian LLC-MK2 cells were obtained from American Type Culture Collection (ATCC; a gift from Nathalie Arhel for the latter); Vero E6 cells were obtained from Sigma-Aldrich (a gift from Christine Chable-Bessia), HEK293T, A549, and MDCK cells were gifts from Michael Malim’s lab and Wendy Barclay’s lab, Huh7 and Huh7.5.1 cells have been described (Nakabayashi et al., 1982; Zhong et al., 2005), respectively, and the latter provided by Raphaël Gaudin. All cell lines were regularly screened for the absence of mycoplasma contamination.

A549 cells (and Caco-2 cells, for the CRISPR screen) stably expressing ACE2 were generated by transduction with RRL.sin.cPPT.SFFV.WPRE containing-vectors (Rebendenne et al., 2021).

For CRISPR-Cas9-mediated gene disruption, Calu-3, Caco-2 and A549-ACE2, cells stably expressing Cas9 or dCas9-VP64 were first generated by transduction with LX_311-Cas9 or XPR_BRD109, respectively, followed by blasticidin selection at 10 µg/ml. WT Cas9 activity was checked using the XPR_047 assay (a gift from David Root, Addgene 107145) and was always >80-90%. dCas9-VP64 activity was checked using the pXPR_502 vector expressing sgRNA targeting IFITM3 and MX1 ISG promoters. Cells were transduced with guide RNA expressing LentiGuide-Puro or XPR_502 (as indicated) and selected with antibiotics for at least 10 days.

### Lentiviral production and transduction

Lentiviral vector stocks were obtained by polyethylenimine (PEI; for LentiGuide vectors) or Lipofectamine 3000 (Thermo Scientific; for XPR_502 vectors)-mediated multiple transfections of 293T cells in 6-well plates with vectors expressing Gag-Pol, the miniviral genome, the Env glycoprotein at a ratio of 1:1:0.5. The culture medium was changed 6 h post-transfection, and vector containing supernatants harvested 36 h later, filtered and used directly or stored at -80**°**C. Transduction was performed by cell incubation with the LV in the presence of polybrene (4 µg/mL) for a few hours. For LX_311-Cas9, XPR_BRD109, RRL.sin.cPPT.SFFV/ACE2.WPRE, LentiGuide-Puro and XPR_502 transductions, spin infection was performed for 2 h at 30°C and 1000g to improve transduction efficiencies.

### CRISPR KO screens

Vero E6, Caco-2-ACE2 and Calu-3 cells were spin infected for 2h at 1000g with LX_311-Cas9 lentiviral vector at a high MOI and in the presence of polybrene (4 µg/mL). Blasticidin selection was added 24-48h post transduction. Cells were grown to at least 120 million cells (40-60 millions for the Calu-3) and transduced with lentiviral vectors coding the C. *sabeus* sgRNAs (Wei et al., 2021) (for Vero E6), the Brunello library (Sanson et al., 2018) (for Caco-2-ACE2) or the Gattinara library (DeWeirdt et al., 2020) (for Calu-3), at MOI ∼0.3-0.5. Puromycin selection was added 24-48h post transduction and maintained for 10-15 days prior to proceeding to the screens. Cells were re-amplified to at least the starting amounts prior to SARS-CoV-2 challenge at MOI 0.005. The day of the viral challenge, 40 million cells were harvested, pelleted by centrifugation and frozen down for subsequent gDNA extraction. Massive CPEs were observed 3-5 days post SARS-CoV-2 infection and cells were kept in culture for 11-13, 18-27, and 30-34 days in total prior to harvest and gDNA extraction, for Vero E6, Caco-2-ACE2 and Calu-3, respectively.

### CRISPRa screens

Calu-3 cells were spin infected for 2h at 1000g with dCas9-VP64 (pXPR_BRD109)- expressing lentiviral vectors at high MOI and in the presence of polybrene (4 ug/mL). Blasticidin selection was added 24-48h post transduction and the cells were amplified. 120 million Calu-3-dCas9-VP64 cells were then transduced with the Calabrese library in two biological replicates (for sublibrary A) or in one replicate (for sublibrary B) at a low MOI (∼0.3-0.5). 2.5 weeks later, 40 million cells were either challenged with SARS-CoV-2 (MOI 0.005) or harvested and frozen down for subsequent gDNA extraction. Massive CPEs were observed 3-5 days post SARS-CoV-2 infection and cells were kept in culture for 11-17 days prior to harvest and gDNA extraction.

### Genomic DNA preparation and sequencing

Genomic DNA (gDNA) was isolated using either the QIAamp DNA Blood Maxi kit (Qiagen) or the NucleoSpin Blood XL kit (Macherey-Nagel), as per the manufacturer’s instructions. Isolated gDNAs were further prepared and cleaned up using a OneStep™ PCR Inhibitor Removal Kit according to manufacturer instructions (Zymo Research, D6030).

### Genomic DNA sequencing

For PCR amplification, gDNA was divided into 100 μL reactions such that each well had at most 10 μg of gDNA. Plasmid DNA (pDNA) was also included at a maximum of 100 pg per well. Per 96 well plate, a master mix consisted of 150 μL DNA Polymerase (Titanium Taq; Takara), 1 mL of 10x buffer, 800 μL of dNTPs (Takara), 50 μL of P5 stagger primer mix (stock at 100 μM concentration), 500 μL of DMSO, and water to bring the final volume to 4 mL. Each well consisted of 50 μL gDNA plus water, 40 μL PCR master mix, and 10 μL of a uniquely barcoded P7 primer (stock at 5 μM concentration). PCR cycling conditions were as follows: an initial 1 min at 95 °C; followed by 30 s at 94 °C, 30 s at 52.5 °C, 30 s at 72 °C, for 28 cycles; and a final 10 min extension at 72 °C. PCR primers were synthesized at Integrated DNA Technologies (IDT). PCR products were purified with Agencourt AMPure XP SPRI beads according to manufacturer’s instructions (Beckman Coulter, A63880). Prior to sequencing the sample was quantitated by qPCR and diluted to 2nM. 5 µL of the sample was then further diluted and denatured with 5 µL 0.1N NaOH and 490 µL HT1 buffer (Illumina). Samples were sequenced on a HiSeq2500 HighOutput (Illumina) with a 5% spike-in of PhiX.

### Screen analysis

For each published screen, corresponding authors provided raw read counts. For the screens conducted in this paper, guide-level read counts were retrieved from sequencing data. We log-normalized read counts using the following formula:

When applicable, we averaged lognorm values across conditions (Poirier, Daelemans, Sanjana). We calculated log-fold changes for each condition relative to pDNA lognorm values. If pDNA reads were not provided for the given screen, pDNA reads from a different screen that used the same library were used (Puschnik analysis used Sanjana pDNA, Zhang analysis used Poirier pDNA). Log-fold changes were used to calculate the receiver-operator characteristic area under the curve values (ROC-AUC) for control populations, where essential genes were treated as true positives and non-essential genes were treated as true negatives. We define essential genes based on Hart et al. 2015 and non-essential genes based on Hart et al. 2014. For each condition in each dataset, we fit a natural cubic spline between the control and infected conditions (Wei et al. 2021). The degrees of freedom for each spline were fit using 10-fold cross-validation. We calculated residuals from this spline and z-scored these values at the guide-level (anchors package). We calculated gene-level z-scores by averaging across guides and conditions, and p-values were combined across conditions using Fisher’s method. Genes were filtered by number of guides per gene, which was generally one guide fewer or greater than the median number of genes per gene for that library (e.g. for Brunello screens, which has a median of 4 guides per gene, we applied a filter of 3 to 5 guides per gene). This guide-filtering step accounts for any missing values in the file compiling data across all screens (all_screens_v3.xlsx). We then used these filtered gene-level z-scores to rank the genes such that the rank one gene corresponded to the top resistance hit. The files containing the guide-level and gene-level residual z-scores for each screen are being deposited on Gene Expression Omnibus (GEO) (**Supplemental Files 1-5**). All code used in this analysis can be found at: https://github.com/PriyankaRoy5/SARS-CoV-2-meta-analysis.

### Wild-type and reporter SARS-CoV-2 production and infection

The (wild-type) BetaCoV/France/IDF0372/2020 isolate was supplied by Pr. Sylvie van der Werf and the National Reference Centre for Respiratory Viruses hosted by Institut Pasteur (Paris, France). The patient sample from which strain BetaCoV/France/IDF0372/2020 was isolated was provided by Dr. X. Lescure and Pr. Y. Yazdanpanah from the Bichat Hospital, Paris, France. The mNeonGreen (mNG) (Xie et al., 2020a) and Nanoluciferase (NLuc) (Xie et al., 2020b) reporter SARS-COV-2 were based on 2019-nCoV/USA_WA1/2020 isolated from the first reported SARS-CoV-2 case in the USA, and provided through World Reference Center for Emerging Viruses and Arboviruses (WRCEVA), and UTMB investigator, Dr. Pei Yong Shi.

WT, mNG and NLuc reporter SARS-CoV-2 were amplified in Vero E6 cells (MOI 0.005) in serum-free media. The supernatant was harvested at 48 h-72 h post infection when cytopathic effects were observed, cell debris were removed by centrifugation, and aliquots frozen down at -80°C. Viral supernatants were titrated by plaque assays in Vero E6 cells. Typical titers were 3.10^6^-3.10^7^ plaque forming units (PFU)/ml.

Simian and human cell infections were performed at the indicated multiplicity of infection (MOI; as calculated from titers in Vero E6 cells) in serum-free DMEM and 5% serum-containing DMEM, respectively. The viral input was left for the duration of the experiment (unless specified otherwise). The viral supernatants were frozen down at -80°C prior to RNA extraction and quantification and/or titration by plaque assays on Vero E6 cells. The cells were trypsinized and the percentage of cells expressing mNG was scored by flow cytometry using a NovoCyte^TM^ (ACEA Biosciences Inc.) after fixation in PBS1X-2% PFA, or the cells were lysed in Passive Lysis buffer and NLuc activity measured inside the BSL-3 facility, or lysed in RLT buffer (Qiagen) followed by RNA extraction and RT-qPCR analysis, at the indicated time post-infection.

### Seasonal coronavirus production and infection

HCoV-229E-Renilla was a gift from Volker Thiel (van den Worm et al., 2012) and was amplified for 5-7 days at 33°C in Huh7.5.1 cells in 5% FCS-containing DMEM. HCoV-NL63 NR-470 was obtained through BEI Resources, NIAID, NIH and was amplified for 5-7 days at 33°C in LLC-MK2 simian cells, in 2% FCS-containing DMEM. Viral stocks were harvested when cells showed >50% CPEs. Viruses were titrated through TCID_50_ in the cells used for their amplification and typical titers were 1,8.10^9^ TCID_50_/mL and 10^6^ TCID_50_/mL for HCoV-229E-Renilla and HCoV-NL63, respectively. Infections of Calu-3 were performed at MOI 300 for HCoV-229E-Renilla (as measured on Huh7.5.1 cells) and MOI 0.1 for HCoV-NL63 (as measured on LLC-MK2 cells) and infection efficiency was analyzed 3 days later by measuring Renilla activity or 5 days later by RT-qPCR for HCoV-229E-Renilla and HCoV-NL63, respectively.

### MERS-CoV production and infection

To produce MERS-CoV, HEK-293T cells were transfected with a bacmid containing a full-length cDNA clone of the MERS-CoV genome (a king gift of Dr Luis Enjuanes; (Almazán et al., 2013)) and overlaid six hours later with Huh7 cells. After lysis of Huh7 cells, cell supernatants were collected and the virus was further amplified on Huh7 cells. Viral stocks were aliquoted and frozen down, and titrated by the TCID_50_ method.

Calu-3 cells, seeded in 24-wells on glass coverslips (immunofluorescence) and in duplicate in 48-wells (infectivity titrations), were inoculated with MERS-CoV at an MOI of 0.3. Sixteen hours after inoculation, coverslips were fixed by incubation in 3% paraformaldehyde for 20 minutes, and stored in PBS at 4°C until immunolabeling was performed. Supernatant was collected from the infected cells in the 48-wells and stored at -80°C until infectivity titrations were performed. Coverslips were further processed for immunolabeling of the infected cells. Briefly, cells were permeabilized by incubation with 0.4% Triton X-100 for 5 minutes, and were then blocked by incubation for 30 minutes with 5% goat serum (GS) in PBS. Infected cells were labelled with a mixture of the mouse monoclonal antibody J2 against dsRNA (Scicons, diluted 1:400) and a rabbit polyclonal antibody directed against the spike protein (Sino Biological Inc, diluted 1:500) in PBS supplemented with 5% GS for 30 minutes at room temperature. They were washed three times with PBS and then incubated for 30 minutes with Alexa-488-conjugated donkey anti-mouse IgG and Alexa594-conjugated goat anti-rabbit IgG secondary antibodies (both from Jackson Immunoresearch) in 5% GS in PBS supplemented with 1 μg/ml DAPI (4′,6- diamidino-2-phenylindole). Coverslips were then rinsed four times with PBS, once in MilliQ water and mounted on microscope slides in Mowiol 4-88-containing medium. Images were acquired on an Evos M5000 imaging system (Thermo Fisher Scientific) equipped with light cubes for DAPI, GFP and TX-RED, and a 10x objective. For each coverslip, ten 8-bit images of each channel were acquired. The total number of cells was determined by counting the nuclei. Infected cells, defined as positive for dsRNA or spike immunolabeling, were counted, and the percentage of infected cells was calculated. About 10,000 to 20,000 cells were counted per condition in each experiment using homemade macros running in ImageJ. For the infectivity titrations, Huh7 cells, seeded in 96 well plates, were inoculated with 100 µl of 1/10 serially diluted supernatants. Cells were incubated with the virus dilutions for 5 days at 37°C. Then, the 50% tissue culture infectious dose (TCID_50_) was determined by assessing the CPEs in each well by light microscopy and the 50% and point was calculated according to the method of Reed and Muench (REED and MUENCH, 1938).

### IAV-NLuc production and infection

The A/Victoria/3/75 virus carrying a NanoLuciferase reporter gene generation and production have been described (Doyle et al., 2018). Viruses were amplified on MDCK cells cultured in serum-free DMEM containing 0.5 μg/mL L-1-p-Tosylamino-2-phenylethyl chloromethyl ketone (TPCK)-treated trypsin (Sigma–Aldrich). Stocks were titrated by plaque assays on MDCK cells.

IAV-NLuc challenges were performed in quadruplicates in 96-well plates, in serum-free DMEM for 1 h and the medium was subsequently replaced with DMEM containing 10% foetal bovine serum. The cells were lysed 10h later and NanoLuc activity was measured with the Nano-Glo assay system (Promega), and luminescence was detected using a plate reader (Infinite 200 PRO; Tecan).

### SARS-CoV-2 internalization assay

Calu-3 cells were incubated with SARS-CoV-2 at an MOI of 5 for 2h at 37°C, were washed twice with PBS and then treated with Subtilisin A (400 μg/mL) in Subtilisin A buffer (Tris/HCl (pH 8.0), 150 mM NaCl, 5 mM CaCl_2_)) in order to get rid of the cell surface-bound viruses, prior to lysis in 350 μL RLT buffer, RNA extraction using the RNeasy kit according the manufacturer’s instructions (Qiagen) and RdRp RT-qPCR to measure the relative amounts of internalized viruses.

### Spike pseudotype production

293T cells were seeded in a 6 well plate prealably coated with poly-Lysine (Sigma-Aldrich) and, 1 day later, transfected with 5 μg of an expression plasmid coding either VSV-G (pMD.G) or SARS-CoV-2 Spike del19 (pcDNA3.1_spike_del19) using Lipofectamine 2000 (Thermo Scientific). The culture medium was replaced after 6h. Cells were infected 24h post-transfection with VSVΔG-GFP-Firefly Luciferase (Rentsch and Zimmer, 2011) at a MOI of 5 for 1h at 37°C and subsequently rinsed 3 times with PBS. The medium was replaced with 5%FCS-supplemented DMEM complemented with a mouse monoclonal anti-VSV-G antibody (CliniSciences, clone 8G5F11, final concentration 1 μg/mL) to neutralize residual viral input, as described (Condor Capcha et al., 2020). Cell supernatants containing pseudotyped VSV viruses were harvested 24h later, spun at 1000 g for 10 min and stored at -80°C.

### Quantification of viral RNAs

3-5 x 10^5^ cells infected or not with SARS-CoV-2 or HCoV-NL63 were harvested and total RNA was extracted using the RNeasy kit (Qiagen) employing on-column DNase treatment, according to the manufacturer’s instructions. 50-125 ng of total RNAs were used to generate cDNAs. To quantify SARS-CoV-2 RNAs, the cDNAs were analyzed by qPCR using published RdRp primers and probe (Corman et al., 2020), as follow: RdRp_for 5’-GTGARATGGTCATGTGTGGCGG-3’, RdRp_rev 5’-CAAATGTTAAAAACACTATTAGCATA-3’, and RdRp_probe 5’-FAM-CAGGTGGAACCTCATCAGGAGATGC-TAMRA-3’). To quantify HCoV-NL63 RNAs, the cDNAs were analyzed by qPCR using published primers and probe (Carbajo-Lozoya et al., 2012), as follow: NL-63F2 5′-CTTCTGGTGACGCTAGTACAGCTTAT-3′, NL-63R2 5′-AGACGTCGTTGTAGATCCCTAACAT-3′, and NL-63 probe 5′-FAM- CAGGTTGCTTAGTGTCCCATCAGATTCAT-TAMRA-3′ (Carbajo-Lozoya et al., 2012). qPCR reactions were performed in triplicate, in universal PCR master mix using 900 nM of each primer and 250 nM probe or the indicated Taqmans. After 10 min at 95°C, reactions were cycled through 15 s at 95°C followed by 1 min at 60°C for 40 repeats. Triplicate reactions were run according to the manufacturer’s instructions using a ViiA7 Real Time PCR system (ThermoFisher Scientific). pRdRp and pNL63 (which respectively contains fragments amplified from SARS-CoV-2- and NL63-infected cell RNAs using primers RdRp_for and RdRp_rev, and NL-63F2 and NL-63R2, cloned into pPCR-Blunt II-TOPO) was diluted in 20 ng/ml salmon sperm DNA to generate a standard curve to calculate relative cDNA copy numbers and confirm the assay linearity (detection limit: 10 molecules of RdRp per reaction).

### ACE2 staining using Spike RBD-mFc recombinant protein and flow cytometry analysis

The SARS-CoV-2 Spike RBD sequence used here as a soluble tagged exofacial ligand for ACE2, was obtained from RNA extracted from a patient nasopharyngeal sample collected in Montpellier University hospital during Spring 2020 and a gift from Vincent Foulongne (Veyrenche et al., 2021) (RBD sequence GenBank accession number MT787505.1). The predicted N-terminal signal peptide of the spike protein (amino acid 1-14) was fused to the RBD sequence (amino acid 319-541) and C-terminally tagged with a mouse IgG1 Fc fragment. The RBD-mFc fusion sequence was then cloned into a pCSI vector for expression in mammalian cells, as previously described (Giovannini et al., 2013). The pCSI-SpikeRBD expression vector was transfected in HEK293T cells using the PEIpro® transfection reagent. Cells were washed 6h post transfection and grown for an additional 72-96h in serum-free Optipro medium (Invitrogen) supplemented with glutamine and non-essential amino acids. Conditioned medium was then harvested, filtered through 0.45 *μ*m filters and concentrated 100-fold by centrifugation at 3600 rpm at 4°C on 10 kDA cut-off Amicon Ultra-15 concentrators. Samples were aliquoted and stored at -20°C until further use.

For ACE2 labelling, cells were harvested and incubated 20 min at 37°C in FACS buffer (PBS1X-2% BSA) containing a 1/20 dilution of Spike RBD-mFc followed by secondary anti-mouse Alexa-488 incubation and several washes in FACS buffer. Flow cytometry was performed using the NovoCyte^TM^ (ACEA Biosciences Inc.).

### Immunoblot analysis

Cells were lysed in lysis buffer (10 mM TRIS 1M pH7.6, NaCl 150 mM, Triton X100 1%, EDTA 1 mM, deoxycholate 0,1%) supplemented with sample buffer (50 mM Tris-HCl pH 6.8, 2% SDS, 5% glycerol, 100 mM DTT, 0.02% bromophenol blue), resolved by SDS-PAGE and analyzed by immunoblotting using primary antibodies against ACE2 (ProteinTech 21115-1-P) and Actin (Sigma-Aldrich A1978), followed by HRP-conjugated anti-rabbit or anti-mouse immunoglobulin antibodies and chemiluminescence Clarity or Clarity max substrate (Bio-Rad). A Bio-Rad ChemiDoc imager was used.

### Analysis of scRNAseq data

For scRNaseq analysis, Seurat objects were downloaded from figshare: (https://doi.org/10.6084/m9.figshare.12436517.v2; (Chua et al., 2020)). Cell identities and CRISPR hits were selected and plotted using the DotPlot function in Seurat (Chua et al., 2020).

### Data availability

The datasets generated during and/or analyzed during the current study are being deposited with GEO and are additionally available from the corresponding authors on reasonable request.

### Requests for materials

Requests for material should be addressed to Caroline Goujon or John Doench at the corresponding address above, or to Addgene for the plasmids with an Addgene number.

## Acknowledgements

We are thankful to Sylvie Van de Werf from the French National Reference Centre for Respiratory Viruses (Pasteur Institute) for providing us with SARS-CoV-2 BetaCoV/France/IDF0372/2020 (isolated by Dr. X. Lescure and Pr. Y. Yazdanpanah, Bichat hospital), to the World Reference Center for Emerging Viruses and Arboviruses (WRCEVA) and UTMB investigator, Dr. Pei Yong Shi, for the mNeon Green and NanoLuc reporter SARS-CoV-2; to Volker Thiel for Renilla reporter HCoV-229E; to BEI Resources, NIAID, NIH for providing us with HCoV-NL63. We wish to thank Raphaël Gaudin, Lucile Espert, Véronique Hebmann, Christine Chable-Bessia, Lise Chauveau, Bruno Beaumelle, Nathalie Arhel, Marc Sitbon, Vincent Foulongne, and Raffaele De Francesco for the generous provision of reagents, and we are thankful to María Moriel-Carretero and Laura Picas for helpful discussions. We are grateful to Christine Chable-Bessia and all CEMIPAI BSL-3 facility members for setting up excellent working conditions for SARS-CoV-2 handling.

This work was supported by the French National Research Agency ANR under the ANR-RA COVID program (ANR-20-COV6-0001, CRISPR-TARGET-CoV, to CG), the European Research Council (ERC) under the European Union’s Horizon 2020 research and innovation programme (grant agreement 759226, ANTIViR, to CG), the Institut National de la Santé et de la Recherche Médicale (INSERM) (to CG), the ‘Fondation CNRS’ (to CG & OM), institutional funds from the Centre National de la Recherche Scientifique (CNRS) and Institut des Sciences Biologiques du CNRS (INSB) (to CG and OM), 3-year PhD studentships from the Ministry of Higher Education and Research (to AR, BB, and JM), a 4th year PhD funding from the Fondation pour la Recherche Médicale (to BB), NIAID R21AI157835 (to JGD), the COVID program from CNRS (to L.D.). We acknowledge the imaging facility MRI, member of the national infrastructure France-BioImaging supported by the French National Research Agency (ANR-10-INBS-04) and the CEMIPAI BSL-3 facility.

## Author contributions

A.R., P.R., J.G.D. and C.G. conceived the study; A.R. and C.G. designed the experiments; A.R. and C.G. performed the CRISPR screens; P.R., H.M., P. DW. And J.G.D. performed the computational analysis; A.R., B.B., A.L.C.V., L.D., O.M. and C.G. performed the BSL-2 and BSL-3 experiments; Y.R., M.T., M.A.A. and Y.L. provided technical help; D.G. provided Spike RBD-mFc for ACE2 staining; F.G.G, J.M., M.W. participated in data interpretation. J.G.D. and C.G. provided overall supervision along with S.B. and J.D.; A.R., P.R., J.G.D. and C.G. wrote the manuscript with input from all authors.

---

**Figure S1.**
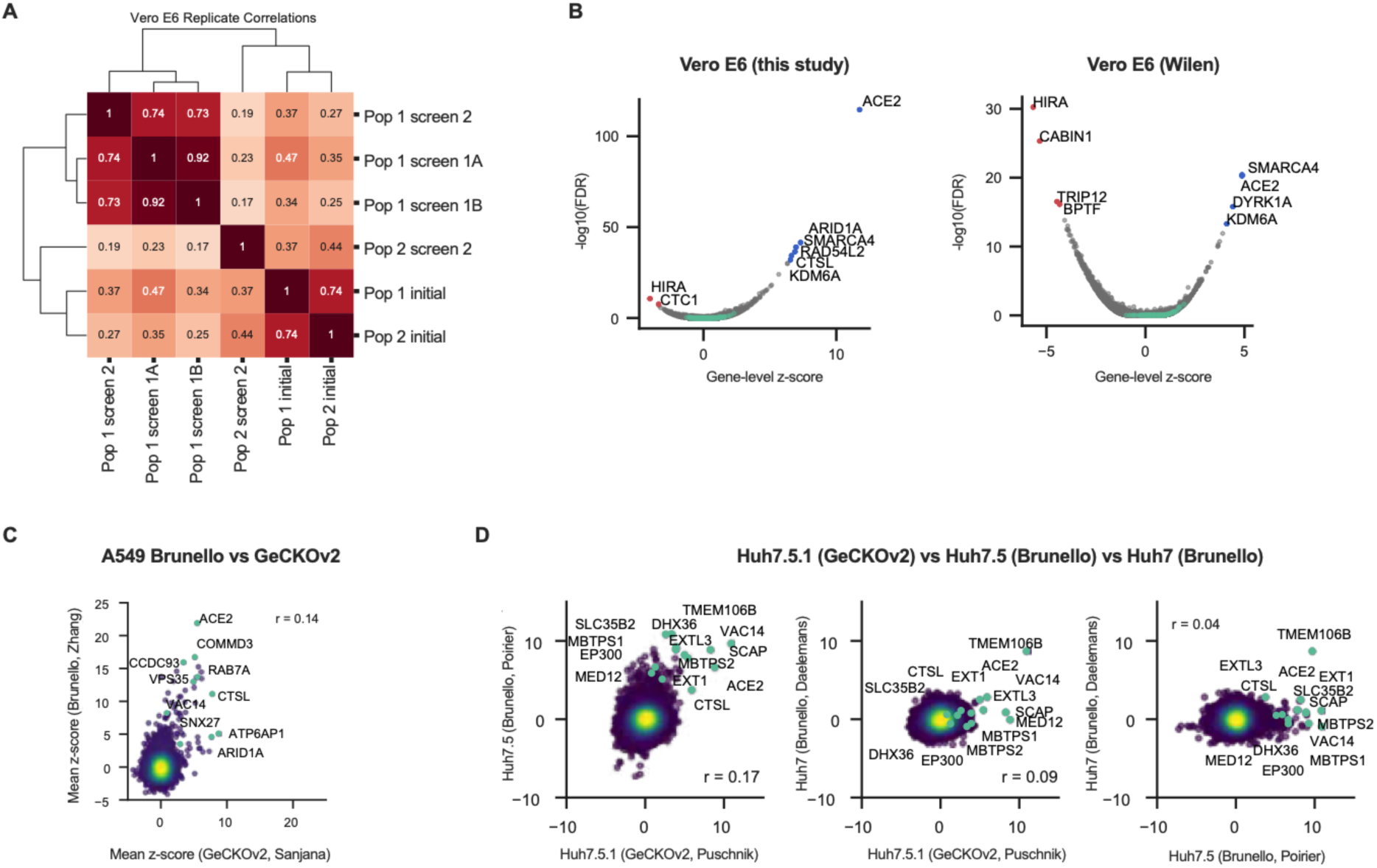
**A.** Clustermap showing correlations of log-fold change values relative to pDNA across replicates in the Vero E6 screen from the present study. Population 1 (Pop 1) and Population 2 (Pop 2) refers to 2 independent library transductions, in which screens 1A, 1B and 2 refer to biological replicates of SARS-CoV-2 infection in Pop 1 and screen 2 refers to one biological replicate of SARS-CoV-2 infection in Pop 2. “Initial” refers to the uninfected condition. **B.** Volcano plot showing the top genes conferring resistance (right, blue) and sensitivity (left, red) to SARS-CoV-2 when knocked out in Vero E6 cells for this screen and the screen conducted by Wei et al. 2021 (Wilen; (Wei et al., 2021)). The gene-level z-score and -log10(FDR) were calculated after averaging across conditions. **C.** Comparison between genome-wide screens conducted in A549 cells overexpressing ACE2 by Daniloski et al. (Sanjana; Daniloski et al., 2021) and Zhu et al. (Zhang; (Zhu et al., 2021)) using the GeCKOv2 and Brunello libraries, respectively. **D.** Pair-wise comparison between genome-wide screens conducted in Huh7.5.1-ACE2, Huh7.5, and Huh7 cells by Wang et al. (Puschnik ; (Wang et al., 2021)), Schneider et al. (Poirier ; (Schneider et al., 2021)), and Baggen et al. (Daelemans ; (Baggen et al., 2021)), respectively. Annotated genes include top 3 resistance hits from each screen as well as genes that scored in multiple cell lines based on the criteria used to construct the Venn diagram in Figure 1D.

**Figure S2.**
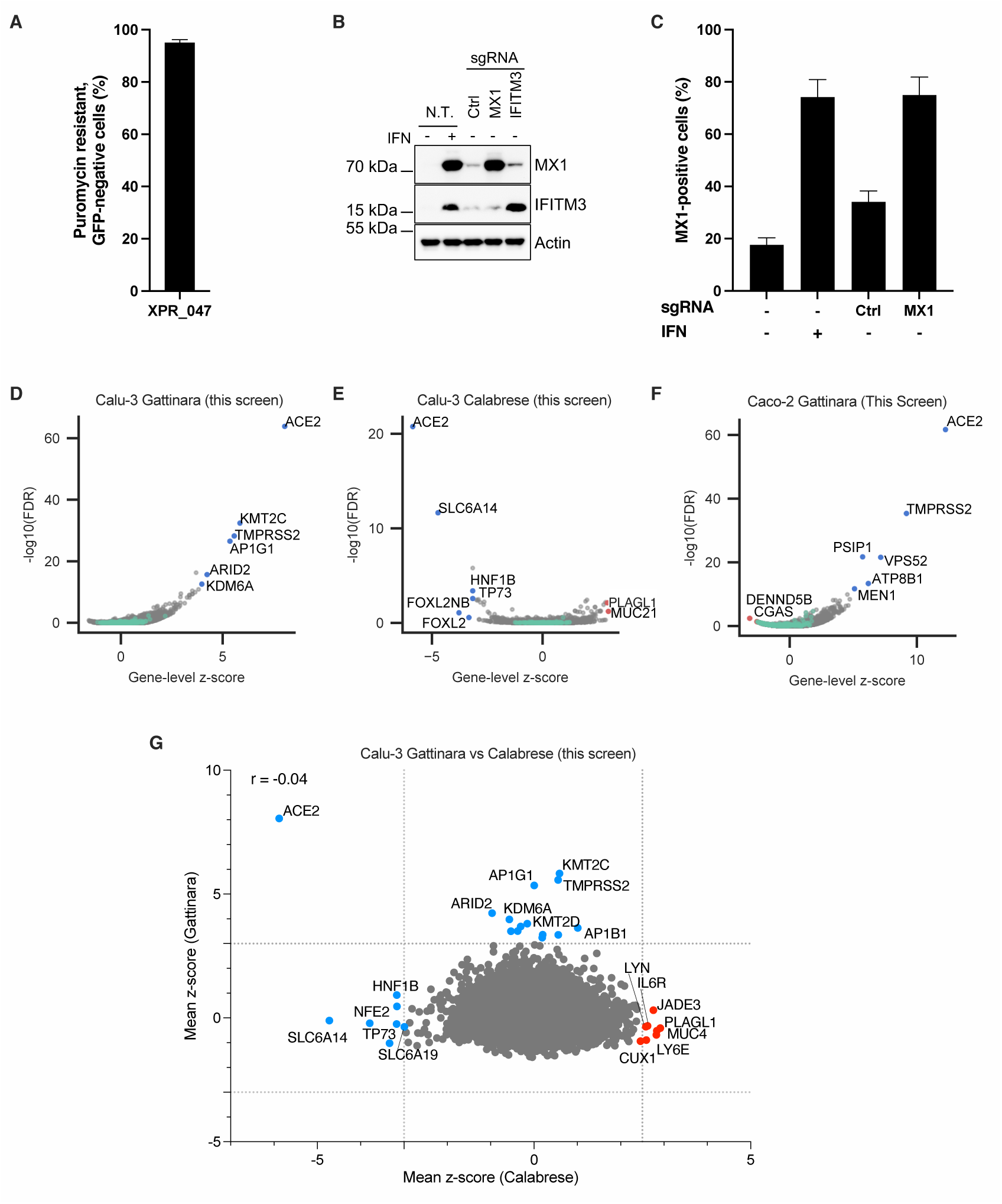
**A.** Calu-3 cells stably expressing Cas9 were transduced with a lentiviral vector expressing the puromycin resistance gene and GFP, as well as a sgRNA targeting the GFP coding sequence (XPR_047). The percentage of puromycin-resistant cells which did not express detectable levels of GFP was scored by flow cytometry 8-10 days post-transduction. **B.** and **C.** Calu-3 cells stably expressing dCas9-VP64 were transduced or not with lentiviral vectors expressing sgRNAs targeting either nothing (Ctrl), *MX1* or *IFITM3* promoter and puromycin-selected for 8-10 days. In parallel, non-transduced (N.T.) cells were treated or not with 1000 U/mL interferon for 24 h. Cells were harvested for immunoblot analysis (**B**) or fixed, permeabilized and stained with an anti-MX1 antibody and an Alexa Fluor 488 secondary antibody and analyzed by flow cytometry (**C**). Biological duplicates (**A, C**) and a representative immunoblot (**B**) are shown. **D.** Volcano plot showing the top genes conferring resistance (right, blue) to SARS-CoV-2 when knocked out in Calu-3 cells. This screen did not have any sensitization hits. The gene-level z-score and -log10(FDR) were calculated after averaging across replicates. **E.** Volcano plot showing the top genes conferring resistance (left, blue) and sensitivity (right, red) to SARS-CoV-2 when overexpressed in Calu-3 cells. The gene-level z-score and -log10(FDR) were calculated after averaging across replicates. **F.** Volcano plot showing the top genes conferring resistance (right, blue) and sensitivity (left, red) to SARS-CoV-2 when knocked out in Caco-2 cells. The gene-level z-score and -log10(FDR) were calculated after averaging across replicates. **G.** Comparison between gene hits in Calu-3 knockout and activation screens. Dotted lines indicated at mean z-scores -3 and 2.5 or 3 for each screen. Proviral and antiviral genes are indicated in blue and red, respectively.

**Figure S3.**
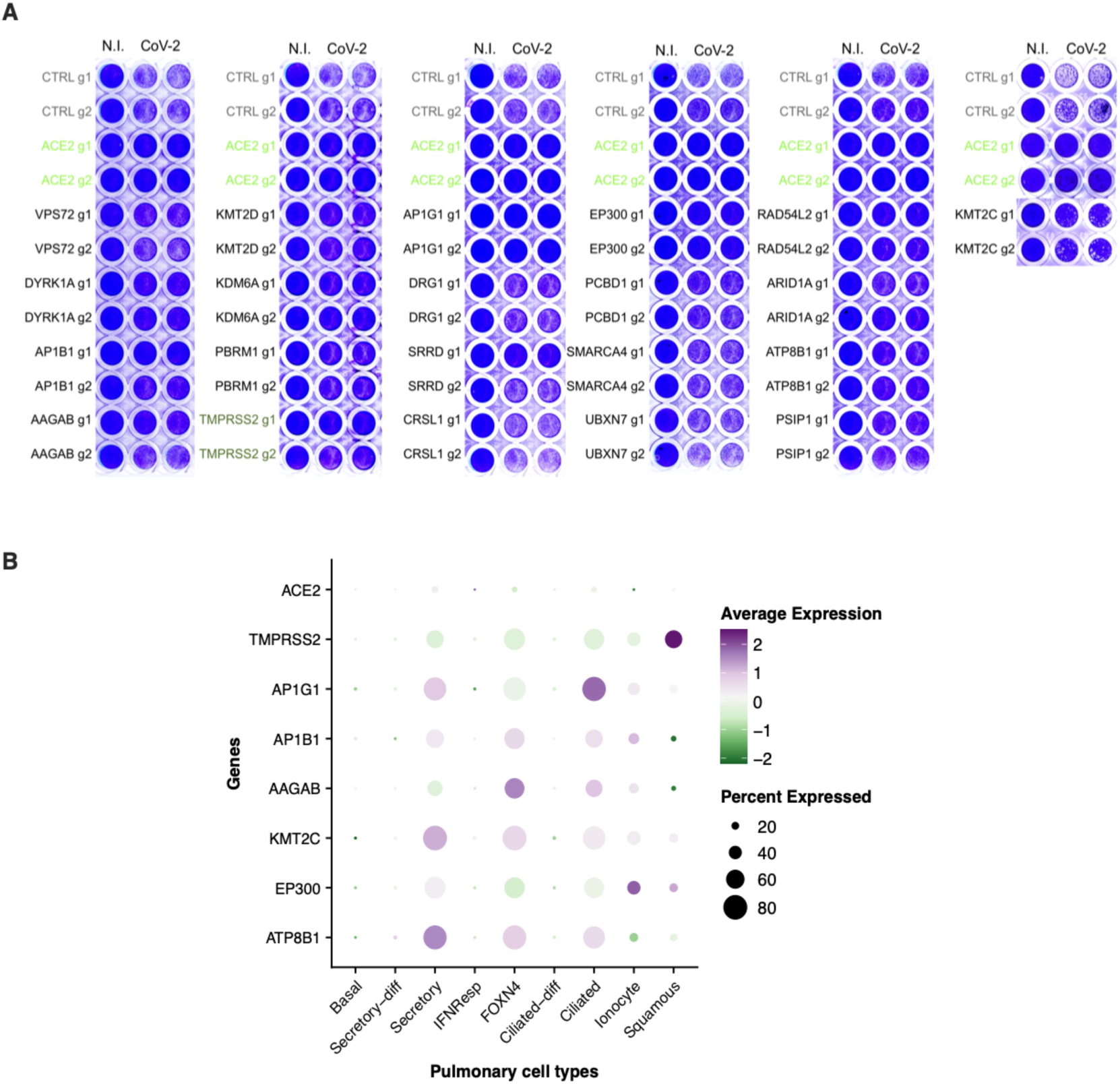
**A. SARS-CoV-2 induced cytopathic effects in candidate KO cell lines.** Calu-3-Cas9 cells were stably transduced to express 2 different sgRNAs (g1, g2) per indicated gene and selected for 10-15 days. Cells were infected by SARS-CoV-2 at MOI 0.005 and ∼5 days later stained with crystal violet. Representative images are shown. **B. Dot plot depicting the expression levels of the best validated genes in the different cell types from the respiratory epithelium**, from Chua et al. data set (Chua et al., 2020). Expression levels in COVID-19 versus healthy patients are color coded; the percentage of cells expressing the respective gene is size coded, as indicated.

**Figure S4.**
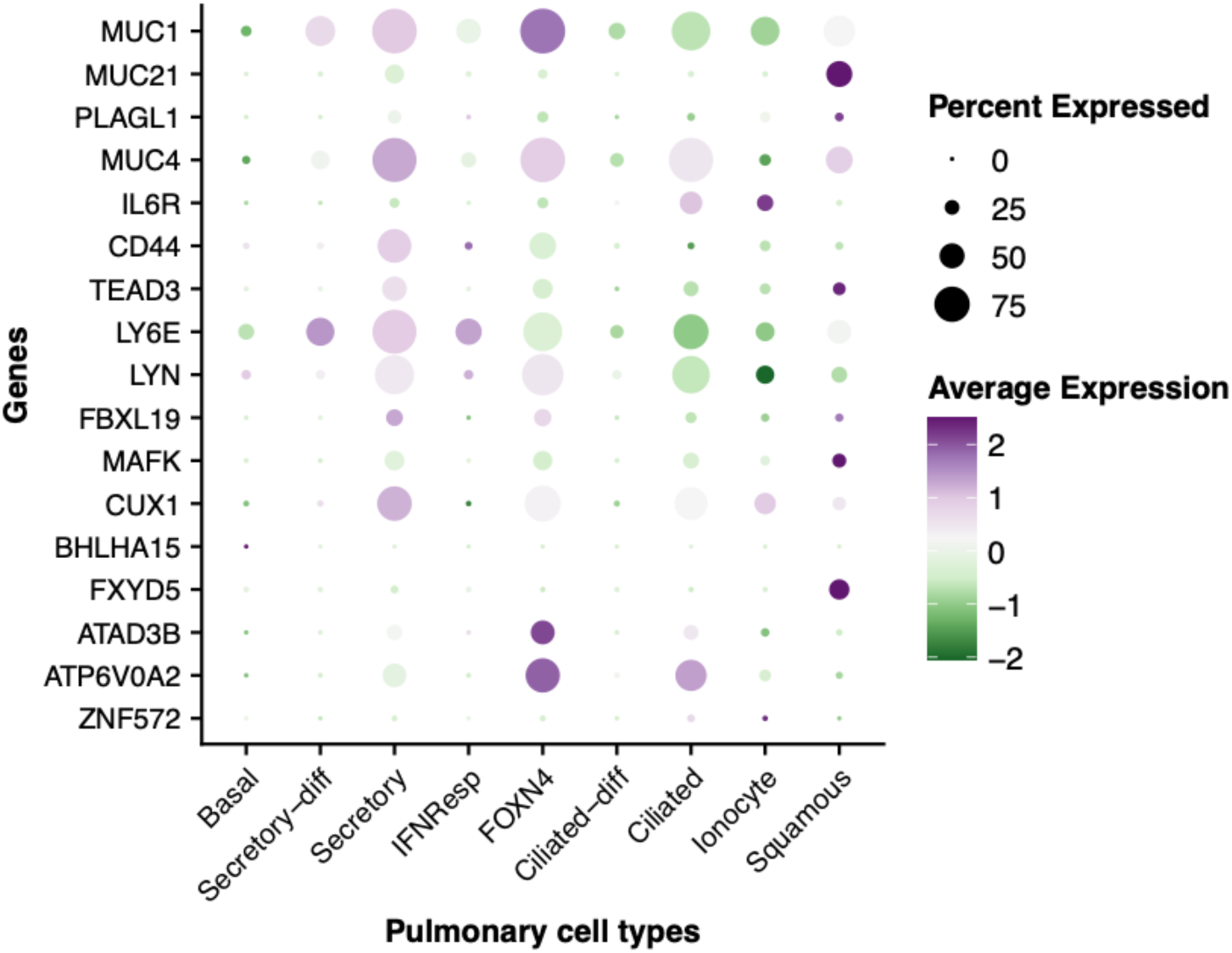
**Dot plot depicting the expression levels of the best validated antiviral genes in the different cell types from the respiratory epithelium**, from Chua et al data set (Chua et al., 2020). Expression levels in COVID-19 versus healthy patients are color coded; the percentage of cells expressing the respective gene is size coded, as indicated.

**Figure S5.**
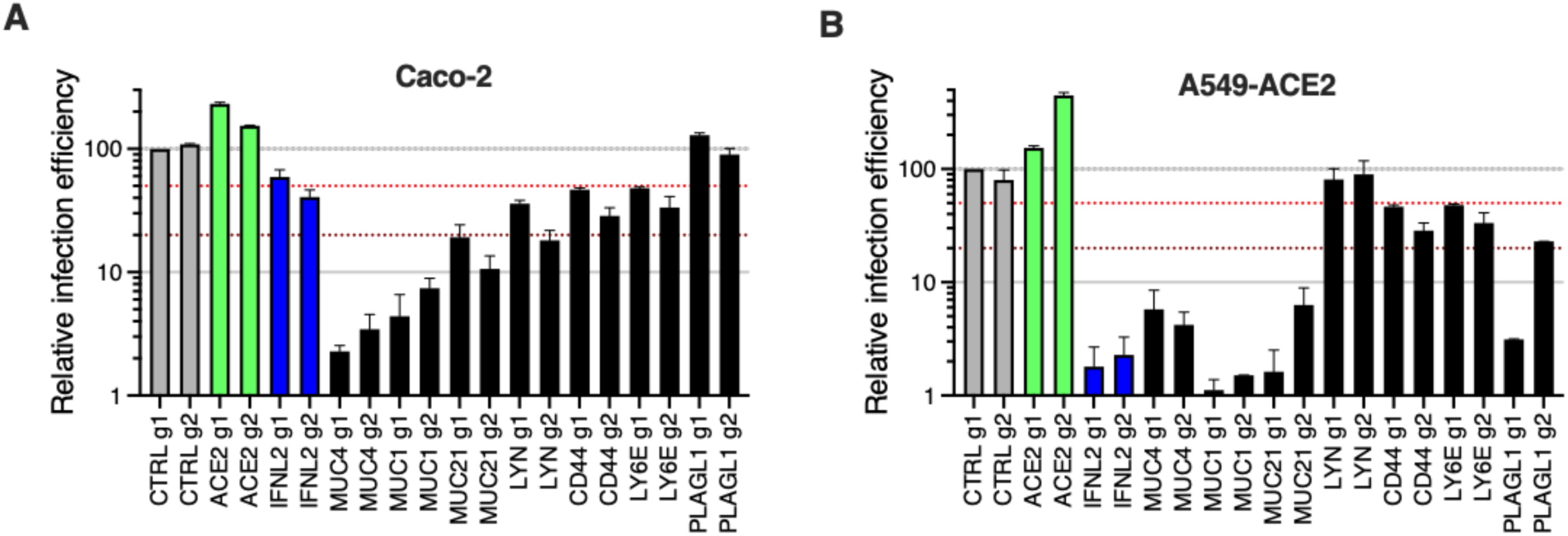
**Impact of the identified antiviral genes on SARS-CoV-2 in Caco-2 and A549-ACE2 cells.** Caco-2-dCas9-VP64 (**A**) and A549-ACE2-dCas9-VP64 (**B**) cells were stably transduced to express 2 different sgRNAs (g1, g2) per indicated gene promoter, or negative controls (CTRL) and selected for at least 10-15 days prior to SARS-CoV-2 mNG infection. The percentage of infected cells was scored 48h later by flow cytometry. Relative infection efficiencies are shown for 2 independent experiments. The red and dark red dashed lines represent 50% and 80% inhibition, respectively.

**Figure S6.**
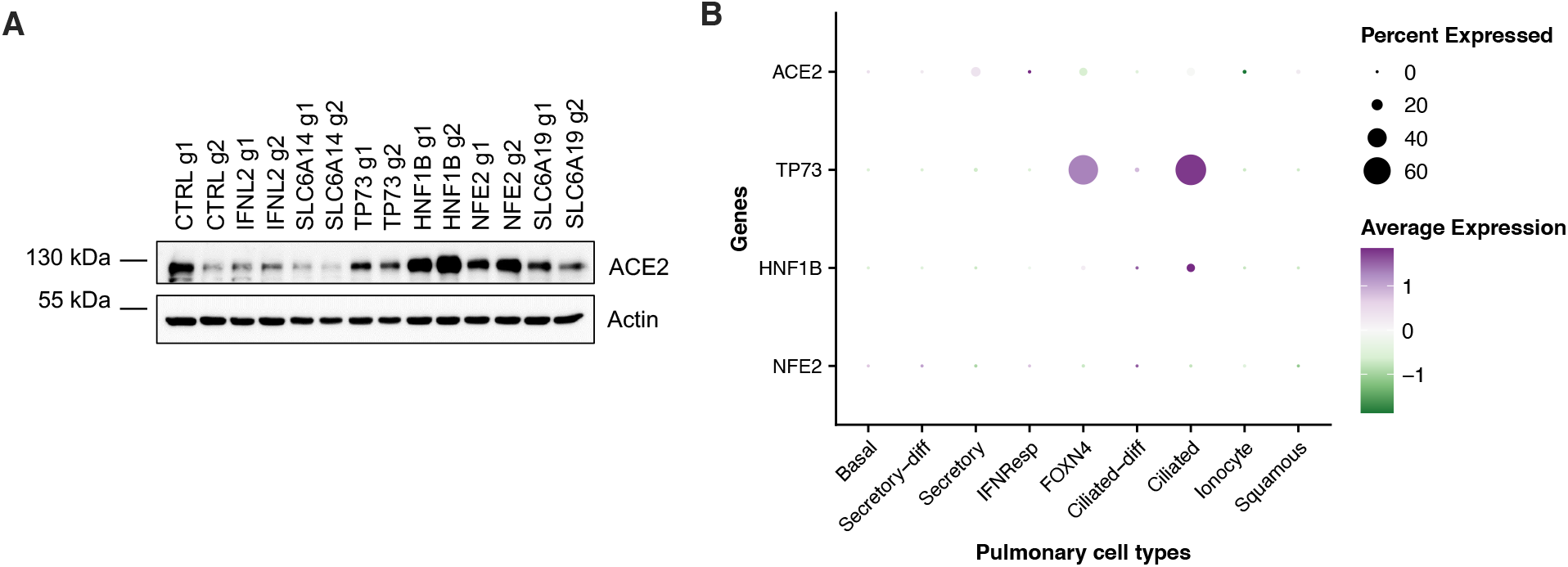
**A. ACE2 expression in CRISPRa cell lines.** Calu-3-Cas9 cells were stably transduced to express 2 different sgRNAs (g1, g2) per indicated gene and selected for 10-15 days (parallel samples from Figures 7-8). The cells were lysed and expression levels of ACE2 were analyzed, Actin served as a loading control. A representative immunoblot is shown. **B. Dot plot depicting the expression levels of the best validated proviral genes in the different cell types from the respiratory epithelium**, from Chua et al data set (Chua et al., 2020). Expression levels in COVID-19 versus healthy patients are color coded; the percentage of cells expressing the respective gene is size coded, as indicated.

**Figure S7.**
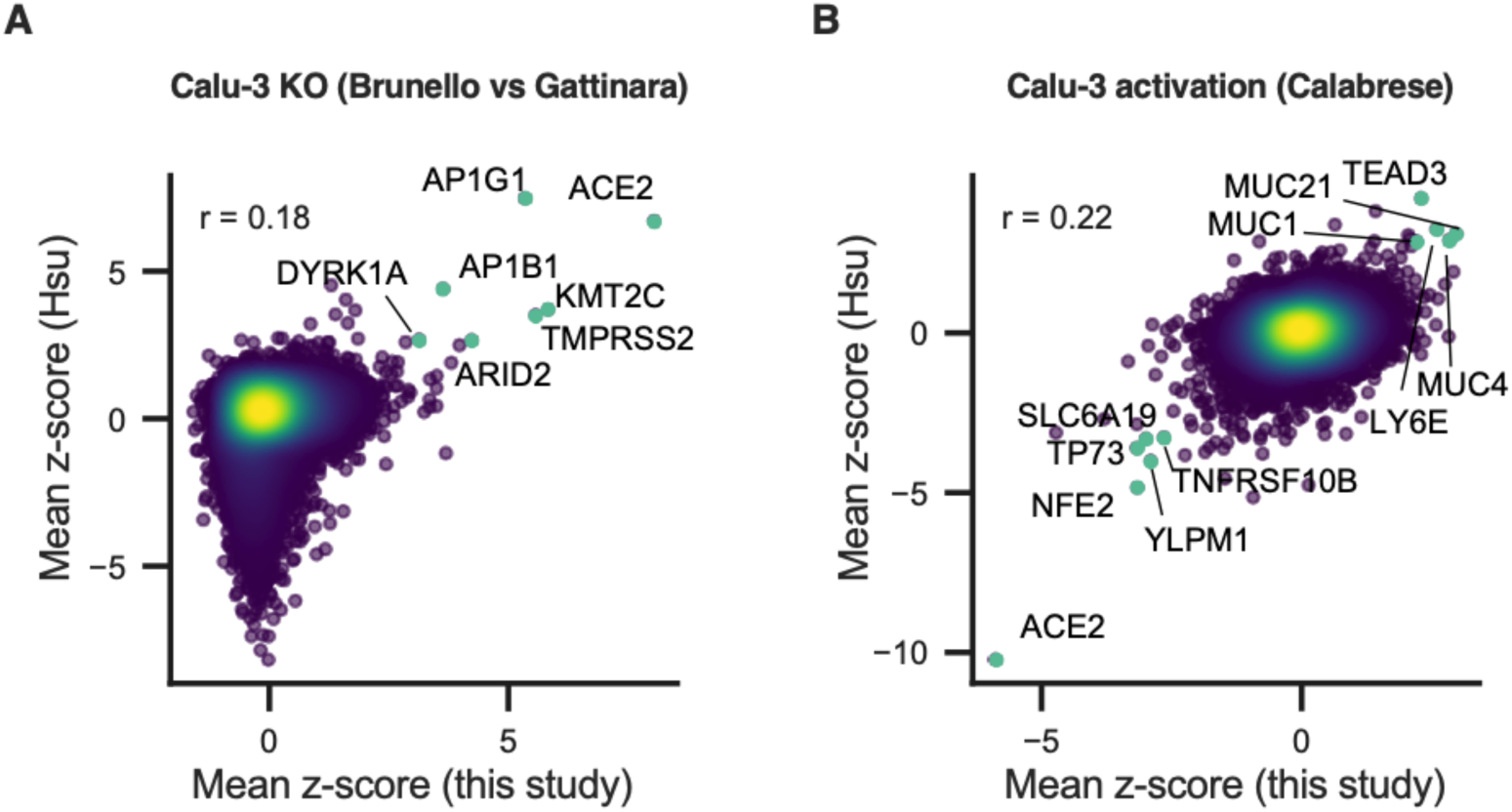
**A.** Comparison between this Calu-3 KO screen to the Calu-3 KO screen conducted by Hsu and colleagues (Biering et al., 2021). Genes that scored among the top 20 resistance hits in both screens are annotated and shown in green. **B.** Comparison between this Calu-3 activation screen to the Calu-3 activation screen conducted by Hsu and colleagues (Biering et al., 2021). Genes that scored among the top 20 resistance hits and sensitization hits in both screens are annotated and shown in green.

